# Blood-brain barrier water exchange measurements using contrast-enhanced ASL

**DOI:** 10.1101/2023.02.16.528529

**Authors:** Elizabeth Powell, Ben R. Dickie, Yolanda Ohene, Geoff J. M. Parker, Laura M. Parkes

**Author notes:** These authors contributed equally to this work (Geoff J. M. Parker, Laura M. Parkes). Corresponding author: **Name** Elizabeth Powell, **Department** Department of Computer Science, **Institute** University College London, **Address** 90 High Holborn, London, WC1V 6LJ, UK, **E-mail**.

## Abstract

A technique for quantifying regional blood-brain barrier (BBB) water exchange rates using contrast-enhanced arterial spin labelling (CE-ASL) is presented and evaluated in simulations and in vivo. The two-compartment ASL model describes the water exchange rate from blood to tissue, *k_b_*, but to estimate *k_b_* in practice it is necessary to separate the intra- and extravascular signals. This is challenging in standard ASL data owing to the small difference in *T*_1_ values. Here, a gadolinium-based contrast agent is used to increase this *T*_1_ difference and enable the signal components to be disentangled. The optimal post-contrast blood *T*_1_ 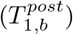 at 3T was determined in a sensitivity analysis, and the accuracy and precision of the method quantified using Monte Carlo simulations. Proof-of-concept data were acquired in six healthy volunteers (five female, age range 24 – 46 years). The sensitivity analysis identified the optimal 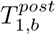 at 3T as 0.8 s. Simulations showed *k_b_* could be estimated in individual cortical regions with a relative error *ϵ* < 1% and coefficient of variation CoV = 30 %; however, a high dependence on blood *T*_1_ was also observed. In volunteer data, mean parameter values in grey matter were: arterial transit time *t_A_* = 1.15±0.49 s, cerebral blood flow *f* = 58.0±14.3 ml blood / min / 100 ml tissue, water exchange rate *k_b_* = 2.32 ± 2.49 s^−1^. CE-ASL can provide regional BBB water exchange rate estimates; however, the clinical utility of the technique is dependent on the achievable accuracy of measured *T*_1_ values.

## Introduction

The blood-brain barrier (BBB) plays a vital role in regulating and maintaining healthy brain function. Passive diffusion of solutes and potential neurotoxins from the blood into the brain is tightly restricted, with the transport of necessary metabolites controlled by specialised proteins. Loss of BBB integrity is increasingly indicated in many neurological conditions, including neurodegeneration (1; 2; 3; 4), stroke (5; 6) and multiple sclerosis (7; 8), as well as more generally in ageing (9; 10; 11). Dynamic contrast-enhanced (DCE) MRI is, at present, the most established MRI method for measuring BBB permeability. When the BBB is damaged, leakage of gadolinium-based contrast agents (GBCA) from blood to brain tissue provides a measurable post-contrast *T*_1_ enhancement. However, DCE-MRI is challenging when BBB damage is subtle as leakage of GBCAs is slow due to the relatively large size of the chelates. Artifacts intrinsic to the method - for example aliasing, signal drift, Gibbs ringing and motion - which can be tolerated when leakage is high, become a limiting factor in detecting the subsequently smaller signal intensity changes for low levels of leakage (12).

Trans-BBB water exchange is an alternative MRI-based biomarker for BBB integrity that has the potential for increased sensitivity to subtle damage (13). Several methods have been developed for measuring trans-BBB water exchange. Proposed arterial spin labelling (ASL) techniques have aimed to separate the intra- and extravascular signals based on diffusion (14; 15; 16) or magnetisation transfer (17) effects, *T*_2_ properties (11; 18; 19; 20; 21) or spatial location (22; 23). Other methods have utilised differences in the *T*_1_ relaxation time (3; 4; 24) or intrinsic diffusion properties (25; 26) of the blood water directly. However, many of the techniques are limited either by long scan times or by a lack of regional exchange rate estimates.

ASL-based methods utilising *T*_1_ differences to separate the intra- and extravascular signals are a potential alternative to the above approaches, and preliminary works manipulating the intravascular *T*_1_ using a GBCA (5; 27)(***citations redacted for double-blind review***) have shown promise. In the absence of GBCAs, measurements of water exchange using this approach are imprecise owing to the small difference in *T*_1_ relaxation times between compartments relative to the exchange rate (28), requiring SNR levels in excess of clinically-attainable values (29). ASL data acquired under the influence of an intravascular GBCA benefit from a larger difference between the intra- and extravascular *T*_1_, which should therefore enable the label location to be determined at lower SNR levels.

Contrast-enhanced (CE) ASL is presented here as a technique for quantifying BBB water exchange, building on previous preliminary data (***citations redacted for double-blind review***). Simulations are first used to determine the optimal post-contrast blood *T*_1_, and then to evaluate the expected accuracy and precision of parameter estimates. Proof-of-concept is then demonstrated in six healthy volunteers.

## Theory

The two-compartment water exchange model (28) for continuous ASL (CASL) describes the imaging voxel in terms of a blood water compartment and an extravascular tissue water compartment, each with corresponding volumes (*v_bw_*, *v_ew_*) and relaxation times (*T*_1,*b*_, *T*_1,*e*_). Following labelling, tagged blood water arrives at the voxel at arterial transit time (ATT), *t_A_*, with a cerebral blood flow (CBF) rate, *f*. Labelled water remains in the intravascular compartment for a finite duration before exchanging into the extravascular compartment. Figure 1A shows a schematic of the compartmental model.

**Figure 1.**
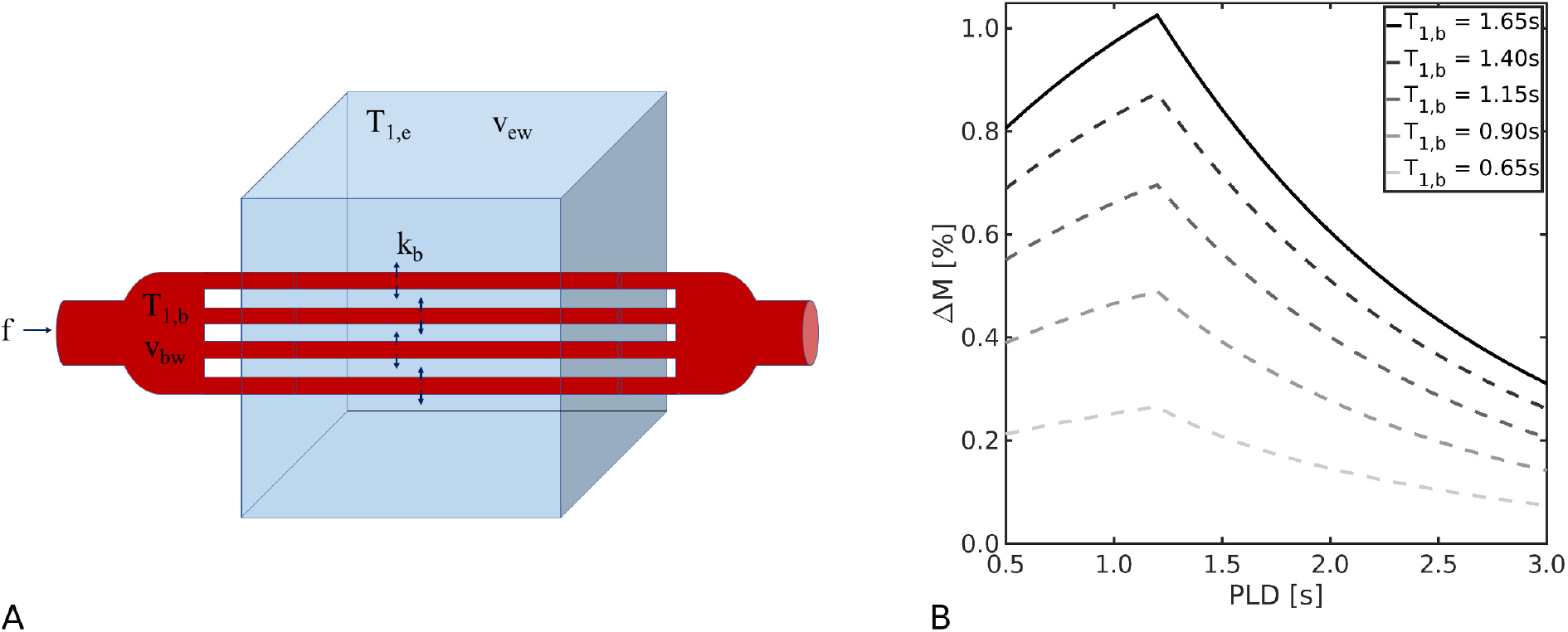
ASL signal model. **(A)**. Schematic diagram of the 2-compartment exchange model (parameter definitions in Table 1). **(B)**. Simulated ASL difference signal Δ*M* for the 2-compartment CASL model in Equation 4 for the equilibrium pre-contrast *T*_1,*b*_ (solid black line) and for a range of post-contrast *T*_1,*b*_ values (dashed lines). Fixed parameters were: exchange rate *k_b_* = 2.65 s^−1^, extravascular relaxation time *T*_1,*e*_ = 1.5 s, cerebral blood flow *f* = 60 ml blood / min / 100ml tissue, label duration *t_L_ = 2* s, arterial transit time *t_A_* = 1.2 s, brain:blood partition coefficient λ = 0.9 and inversion efficiency of labelling *α* = 0.85.

Evolution of each compartment’s magnetisation in the ASL difference image (control - label) is given by:

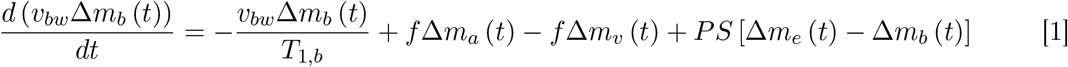

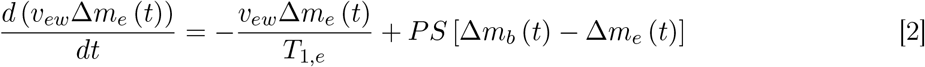

where *PS* is the permeability (*P*) surface area (*S*) product describing exchange between compartments, Δ*m_b_* and Δ*m_e_* represent the magnetisation of capillary blood water and extravascular water within the tissue voxel, and Δ*m_a_* and Δ*m_v_* represent the magnetisation of arterial blood water and venous blood water arriving at and leaving from the tissue voxel respectively. The total ASL difference signal is modelled as the sum of the intra- and extravascular difference magnetisations weighted by their relative volumes:

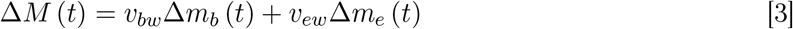

Implicit in Equations 1 and 2 is the assumption that labelled blood resides in exchanging vessels (capillaries and arterioles) only, meaning that contributions from larger vessels (arteries) are excluded; this is generally expected to be valid for post-labelling delay (PLD) times greater than 1 s. Further assumptions can be made to simplify the solutions under certain conditions (28). Firstly, for perfusion rates in normal human brain tissue, it can be assumed that the label will have decayed (due to *T*_1_ recovery) before entering the venous circulation, meaning there is no outflow of labelled blood from the voxel during the PLD and so the venous component can be excluded (i.e. Δ*m_v_* = 0). Secondly, effects of backflow on the signal - that is, exchange of labelled magnetisation from the extravascular space back into the blood - can also be neglected under the assumption that at all times the proportion of labelled extravascular spins is much less than the proportion of labelled intravascular spins (i.e. Δ*m_e_* ≪ Δ*m_b_*, giving PSΔ*m_e_* = 0).

Under these assumptions, as derived in earlier work (28), the time-dependent solutions to Equations 1 and 2 are:

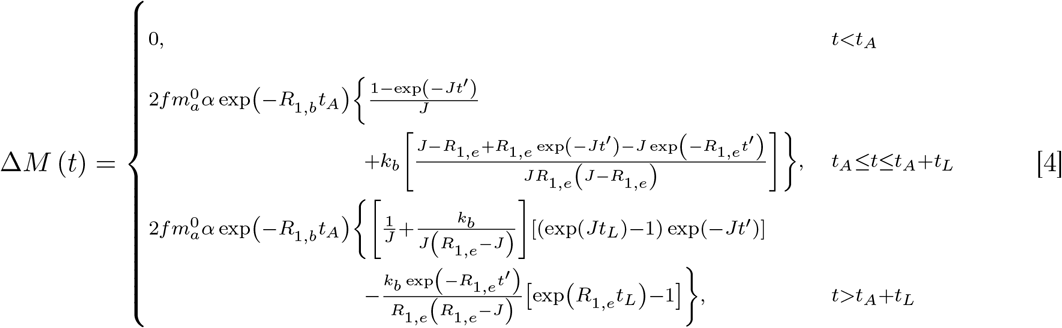

where *t* is the time from the start of labelling, *t_L_* is the labelling duration (LD), *t_A_* is the ATT, 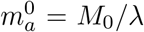 is the equilibrium arterial magnetisation with *M*_0_ the equilibrium magnetisation and λ the brain:blood partition coefficient, *α* is the inversion efficiency of the labelling, *R*_1,*b*_ = ^1^/_*T*_1,*b*__ and *R*_1,*e*_ = ^1^/_*T*_1,*e*__ are the relaxation rates of the blood and tissue compartments, *J* = *k_b_* + *R*_1,*b*_ where the exchange rate of labelled water from blood to tissue is *k_b_* = *PS*/*v_bw_*, and *t*′ = *t* – *t_A_*. Table 1 gives a full definition of all parameters.

**Table 1.**
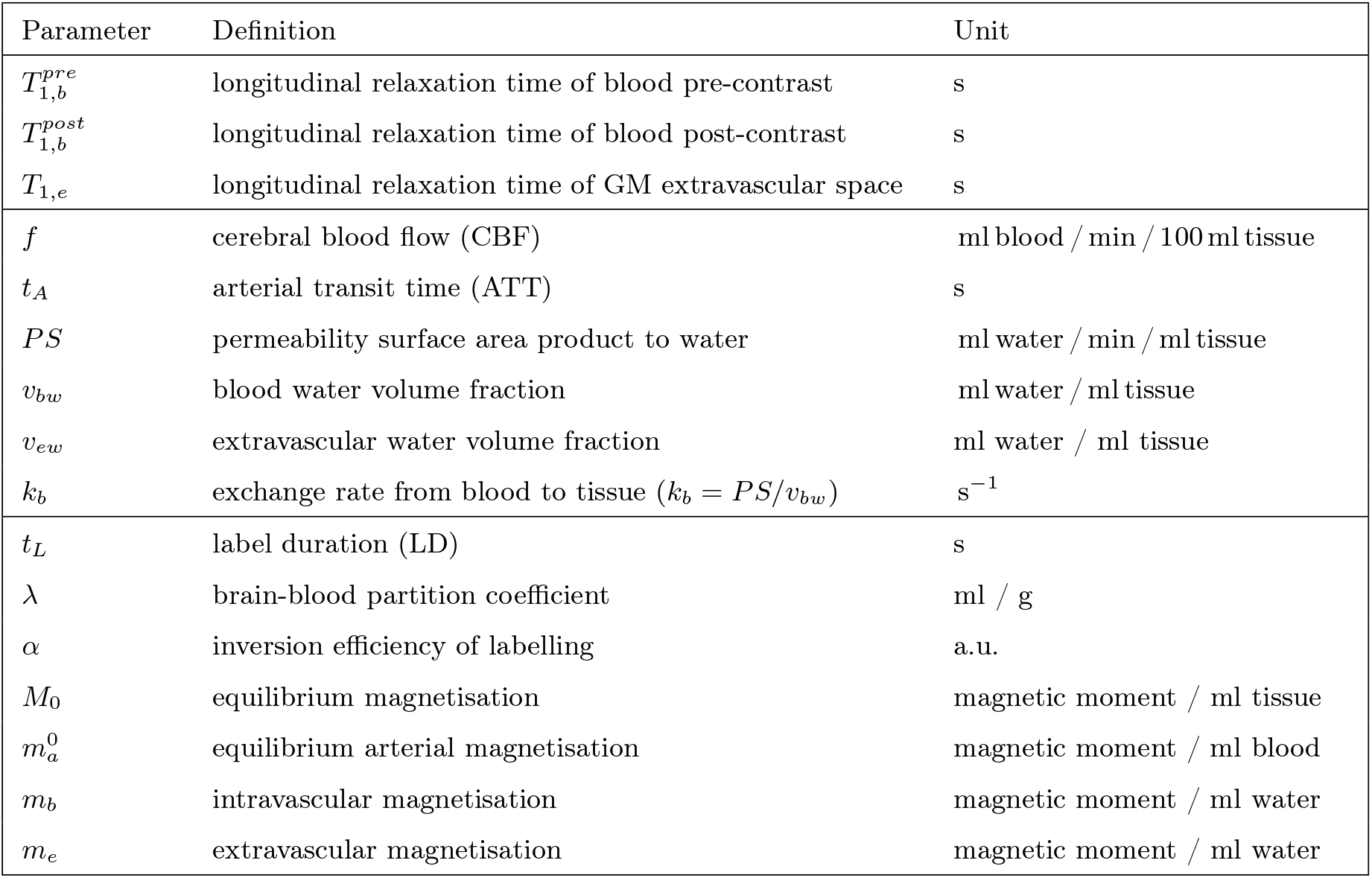
Parameter definitions and units.

| Parameter | Definition | Unit |
| --- | --- | --- |
| $T_{1,b}^{pre}$ | longitudinal relaxation time of blood pre-contrast | s |
| $T_{1,b}^{post}$ | longitudinal relaxation time of blood post-contrast | s |
| $T_{1,e}$ | longitudinal relaxation time of GM extravascular space | s |
| $f$ | cerebral blood flow (CBF) | ml blood / min / 100 ml tissue |
| $t_A$ | arterial transit time (ATT) | s |
| $PS$ | permeability surface area product to water | ml water / min / ml tissue |
| $v_{bw}$ | blood water volume fraction | ml water / ml tissue |
| $v_{ew}$ | extravascular water volume fraction | ml water / ml tissue |
| $k_b$ | exchange rate from blood to tissue ($k_b = PS/v_{bw}$) | $s^{-1}$ |
| $t_L$ | label duration (LD) | s |
| $\lambda$ | brain-blood partition coefficient | ml / g |
| $\alpha$ | inversion efficiency of labelling | a.u. |
| $M_0$ | equilibrium magnetisation | magnetic moment / ml tissue |
| $m_a^0$ | equilibrium arterial magnetisation | magnetic moment / ml blood |
| $m_b$ | intravascular magnetisation | magnetic moment / ml water |
| $m_e$ | extravascular magnetisation | magnetic moment / ml water |

Longitudinal relaxation during the ATT as the labelled blood water arrives at the imaging slice reduces the magnetisation difference Δ*m_b_* (*t*) according to Equation 1. Under the influence of an intravascular GBCA, the shorter blood water *T*_1_ causes Δ*m_b_* (*t*) to reduce more rapidly; this allows the presence of magnetisation that has permeated into the extravascular space (which now has a substantially longer *T*_1_ relative to blood) to have a greater influence on the total difference magnetisation Δ*M* (*t*). Figure 1B displays numerical simulations that illustrate the expected Δ*M* (*t*). With knowledge of the blood and tissue *T*_1_ before and after GBCA contrast injection, Equations 1 and 2 allow these different ΔM (t) to be modelled to extract estimates of the exchange rate. However, as is evident in Figure 1B, higher GBCA concentrations also reduce the overall Δ*M* (*t*), leading to worsened contrast-to-noise ratio. This trade-off is explored here to identify the optimal conditions for CE-ASL estimates of BBB water exchange.

## Methods

Four simulation experiments were performed to assess the feasibility of the CE-ASL method and inform acquisition parameters in vivo. Sensitivity of the CE-ASL signal to water exchange was first evaluated to identify optimal post-contrast blood water *T*_1_ 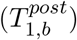 and PLD times at 3 T. Under these optimal conditions, the impact of inaccurate relaxation time values on parameter estimates was explored in an error analysis. Monte Carlo simulations under varying noise conditions then provided an estimate of the expected accuracy and precision of fitted parameters. Finally, the GBCA dose and time post-injection required to obtain optimal 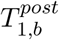 values in vivo were calculated using data from a previous study. Based on the simulation results, an in vivo protocol was designed and conducted in six healthy volunteers. All simulations were performed in Matlab 2019b (The MathWorks).

### Sensitivity analysis

The sensitivity functions were defined as the partial derivative of the signal model in Equation 4 with respect to *k_b_* (provided in full in Appendix A).

To determine optimal 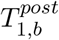 and PLD times, the sensitivity functions were computed for parameter combinations in the ranges 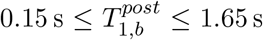 and 0.5 s ≤ PLD ≤ 3.0 s, with *T*_1,*b*_ = 1.65 s taken as the non-contrast-enhanced value in blood at 3T (30). The exchange rate was fixed at *k_b_* = 2.65 s^−1^, the mean of several published studies (13). Calculated sensitivities were normalised using the maximum value obtained across the range of parameter combinations; optimal 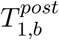 and PLD times were taken as those which maximised the sensitivity functions.

As the modelling approach assumes the GBCA remains intravascular, the impact of extravasated contrast agent on sensitivity to *k_b_* was assessed by varying *T*_1,*e*_ from its non-contrast-enhanced value (1.5 s) (31; 32) down to the optimal 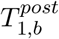 (0.8 s, found previously), thereby mimicking GBCA leakage into tissue and the subsequent reduction of *T*_1,*e*_. For completeness, this range of *T*_1,*e*_ values encompasses the spectrum of exchange rates from no exchange (i.e. *T*_1,*e*_ = 1.5 s) to infinite exchange (i.e. *T*_1,*e*_ = 0.8 s); however, only small reductions are expected for subtle BBB damage. The 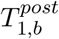 and PLD times were fixed to their optimal values (found previously).

Lastly, variation of the sensitivity in relation to underlying exchange rate was explored for 0.5 s^−1^ ≤ *k_b_* ≤ 4.0 s^−1^, which is representative of previously reported values in human grey matter (GM) (13), over the range 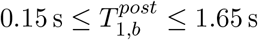.

All other model parameters used in each simulation are provided in Table 2.

**Table 2.**
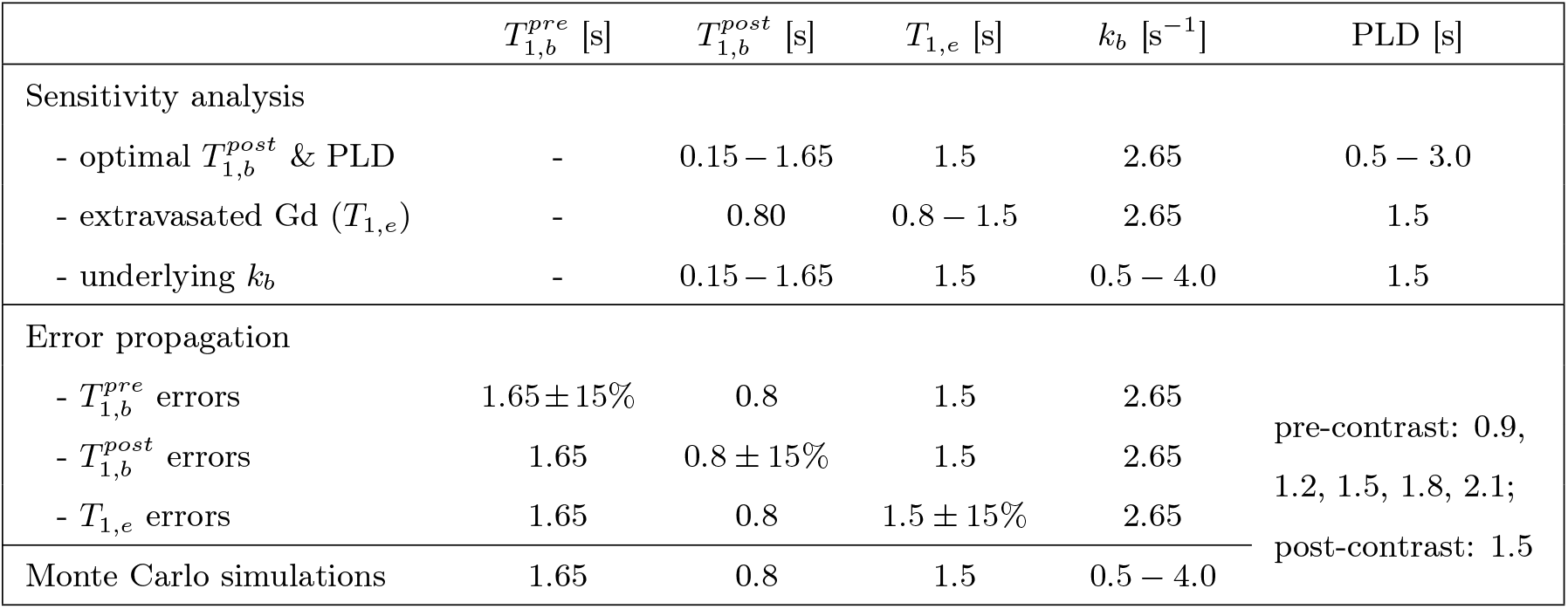
Parameters used in simulations. For all simulations, other fixed parameters were: cerebral blood flow *f* = 60 ml blood / min / 100 ml tissue (29), label duration *t_L_* = 2 s, arterial transit time tA = 1.2 s (30) (note this is variable in vivo depending on labelling location and brain region), brain:blood partition coefficient λ = 0.9 (30) and inversion efficiency *α* = 0.85 (30).

### Error propagation

Systematic biases in parameter estimates arising from *T*_1_ errors were evaluated using numerical simulations. Noise-free synthetic signals were generated using Equation 4 for five PLD times between 0.9 – 2.1 s with *T*_1,*b*_ set at the equilibrium (pre-contrast) value and for one PLD of 1.5 s with *T*_1,*b*_ set at the optimally-reduced (post-contrast) value. Table 2 provides all model parameters. The signal model was then fitted back to the data using perturbations from ground truth *T*_1_ values of ±15%. Fitting was performed using least-squares minimisation with *f*, *t_A_* and *k_b_* as the free parameters initialised using 100 starting values and constrained to 0 ≤ *f* ≤ 200 ml blood / min / 100 ml tissue, 0 ≤ *t_A_* ≤ 2.5 s and 0 ≤ *k_b_* ≤ 5 s^−1^. Starting values were randomly distributed between parameter bounds. Resulting errors in f, *t_A_* and *k_b_* were quantified using the percent relative error *ϵ* = 100 × (*x_fit_* – *x_gt_*)/*x_gt_*, where *x_fit_* and *x_gt_* represent the fitted and ground truth value of a given parameter respectively.

### Accuracy and precision

The accuracy and precision of fitted parameters were estimated using Monte Carlo simulations under varying noise conditions. Data were simulated pre- and post-contrast (as described for the error propagation) for 25 *k_b_* values between 0.5 – 4.0 s^−1^; details of other parameters are in Table 2. For each parameter combination, 2500 control and label signals were synthesised. Zero-mean Gaussian noise with standard deviations *σ* = 0.0033, 0.0017, 0.0011 was added to the control and labelled data independently, giving voxel-wise SNRs of 15, 30, 45 in background-suppressed control data (33) (signal taken as 5% of the equilibrium magnetisation, assuming 95% background suppression efficiency), before pairwise subtraction to create the difference signal. Corresponding voxel-wise SNRs in the difference signal were 1.8, 3.6, 5.4. Voxel-level SNR values were increased by 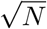 to simulate the higher SNR at regional levels, with *N* = 500 taken as the approximate number of voxels in a cortical region of interest (ROI). All *T*_1_ values were fixed to their ground truth for fitting (performed as for the error propagation). The accuracy of parameter estimates was assessed using the relative error between the ground truth and median fitted values; precision was quantified using the coefficient of variation (CoV), defined as the interquartile range (IQR) of fitted values normalised by the ground truth value. Extreme parameter fits within 5% of the fit constraints (0 ≤ *k_b_* ≤ 10 s^−1^) were discarded from these calculations.

To assess the feasibility of *k_b_* estimates at different regional levels, voxel-wise SNR values were adjusted for signal averaging across ROI sizes equivalent to whole lobes (*N* = 10000) down to the voxel level (*N* = 1) for a single fixed exchange rate of *k_b_* = 2.65 s^−1^ (fitting constrained to 0 ≤ *k_b_* ≤ 5 s^−1^).

### Optimal injected GBCA dose

The GBCA dose needed to achieve the optimal 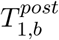 was investigated as a function of injected dose and time post-injection. Using the mean vascular input function (VIF) from the sagittal sinus of 31 healthy volunteers (mean age 66 years, range 52-81 years), 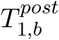 was calculated following 0.1 mmol/kg injection of Dotarem GBCA using image volumes collected every 7.6 s up to 20 min post-injection. These volunteer data were taken from a previous study by Al-Bachari et al. (34).

To estimate the blood concentration (*c_b_*) over time, the functional form of the population-based VIF (35) was fitted to the data:

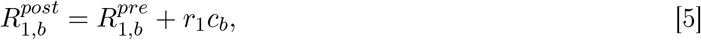

with 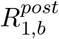 the blood relaxation rate post-contrast, 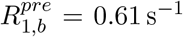 the blood relaxation rate precontrast, and *r*_1_ = 3.4 s^−1^ mM^−1^ the GBCA longitudinal relaxation coefficient (36). Using the estimated blood concentration, the measurements were then scaled to 0.25, 0.50 and 0.75 of the full GBCA dose to simulate 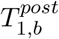 variation at different dose levels. This provided an estimate of the appropriate dose and time post-injection for the optimal 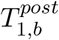.

### MRI acquisition

Proof-of-concept data were acquired in six healthy volunteers (five female, mean age 30 years, range 24 – 46 years) on a simultaneous 3T SIGNA PET-MR scanner (GE Healthcare); ethics approval was granted by the University of Manchester Research Ethics Committee (reference: 2021-5795-18124).

A 3D *T*_1_-weighted MPRAGE image was acquired prior to contrast agent injection with 1mm^3^ isotropic resolution for segmentation of GM, white matter (WM) and CSF.

ASL data and additional *T*_1_ maps were collected pre- and post-contrast agent injection (Figure 2). Two low-dose injections of a GBCA (Dotarem) were administered - each a quarter dose (0.025 mmol/kg), providing 0.050 mmol/kg of Dotarem total - in order to capture the optimal *T*_1_ reduction. Each post-contrast data set (referred to as PC1 and PC2 respectively) was analysed independently.

**Figure 2.**
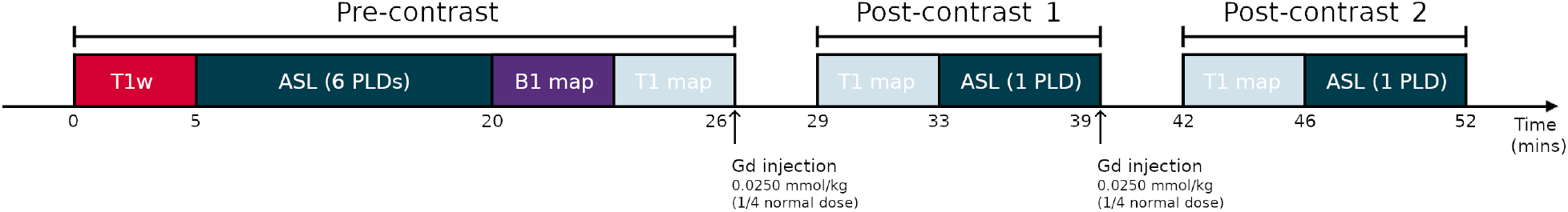
Acquisition pipeline. ASL data and *T*_1_ maps were acquired pre-contrast and again following two low-dose gadolinium injections.

ASL was performed with pseudo-continuous labelling (pCASL), background suppression (all PLDs), no vascular crushing gradients (30), a 3D spiral fast spin echo readout with 8 spiral interleaves, voxel size 1.7 × 1.7 × 4mm^3^ with 36 axial slices covering the complete brain (lowest slice positioned at the base of the cerebellum), TE = 11 ms and minimum TR set according to PLD. The labelling plane was positioned 2 cm inferior and parallel to the 3D acquisition box. Data at six PLDs (0.7, 0.9, 1.2, 1.5, 1.8 and 2.1 s) were collected pre-contrast agent injection (the PLD at 0.7 s was not collected in two subjects), with labelling duration 2 s and 2 repeats (NEX). An additional proton density image was acquired with each PLD. The total pre-contrast acquisition time was 12 minutes. Post-contrast ASL data were collected approximately 7 minutes after each contrast-agent injection at a single PLD of 1.5 s with NEX = 5. One PLD was used to allow time to increase the NEX in comparison to the pre-contrast acquisition and compensate for the expected signal reduction. The acquisition time of each post-contrast data set was 6 minutes.

To produce *T*_1_ maps pre- and post-contrast, 3D *T*_1_-weighted spoiled gradient echo (SPGR) images were acquired using 4 flip angles (2 °, 5 °, 15 ° and 20 °), with voxel size 2 × 2 × 4mm^3^, TR/TE = 4.75/1.06 ms and NEX = 8. Acquisition of the 4 different flip angle images were repeated approximately 3 minutes after each contrast agent injection, prior to the ASL acquisition. Each flip angle acquisition was 1 minute. A 2D Bloch-Siegert B_1_ map was also collected pre-contrast with flip angle 10 °, FOV matched to the *T*_1_ map and resolution 3 × 3 × 8mm^3^.

### MRI analysis

#### Extraction of regional ASL and tissue *T*_1_ values

The ASL subtraction images were divided by the ASL proton density images on a voxel-wise basis. The 3D *T*_1_-weighted image was segmented into GM, WM and CSF using SPM12 (37). Pre- and post-contrast *T*_1_ maps and the ASL proton density image were co-registered to the 3D *T*_1_-weighted image, and the transformation used to propagate the ASL subtraction images into the same space. The *T*_1_-weighted image was then registered to the MNI template, and the transformation applied to the GM and CSF probability maps from the segmentation and the co-registered *T*_1_ maps and ASL subtraction images. The automatic anatomical labelling atlas (38) (masked for GM) was used to extract the mean ASL subtraction signal and *T*_1_ estimates from the 90 cortical and sub-cortical regions (excluding the cerebellum).

#### Estimation of blood *T*_1_

Pre- and post-contrast *T*_1_ maps were estimated by fitting the SPGR signal model to the 4 flip angles at each contrast level (39). This was done in R (version 4.2) using the Levenberg-Marquardt optimisation solver. Each post-contrast *T*_1_ map (in MNI space) was subtracted from the pre-contrast map to produce two subtraction images, one for each contrast agent dose. On each subtraction image a region in the sagittal sinus and straight sinus was identified using the ROI tool in MRIcro (40). The region was masked to contain only those voxels with at least at 20% reduction in *T*_1_ following contrast agent injection; effectively identifying the voxels with the highest blood volume, producing the final blood ROI. The 75th percentile *T*_1_ value within the blood ROI on the pre-contrast *T*_1_ map was recorded as the 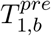 value, chosen as it likely contains high blood volume but will be less affected by noise than the maximum value. The 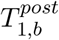 value was estimated by subtracting the prepost *T*_1_ difference (again taken as the 75th percentile value within the blood ROI on the subtraction image) from the 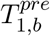 value. This process was repeated independently for each subtraction image.

#### Kinetic modelling of ASL data

Equation 4 was fitted to the data on a voxel-wise basis in Matlab 2021a using an unconstrained simplex search method with initial values *f* = 60 ml blood / min / 100 ml tissue, *t_A_* = 1.0 s, *k_b_* = 1 s^−1^. Voxel-wise *T*_1,*e*_ values and global *T*_1,*b*_ values were fixed to their measured values pre- and post-contrast, with *α* = 0.85 and λ = 0.9. Regional parameter estimates were obtained by taking the median of voxel-wise values within an ROI.

#### SNR estimation

No independent noise measurement was available for the in vivo data, so voxel-level SNR was approximated within each ROI (after averaging signal repetitions) as:

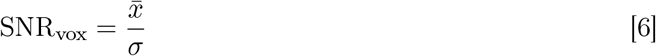

where 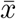 and *σ* are the signal mean and SD within an ROI in the ASL difference image at PLD = 1.5 s.

## Results

### Sensitivity analysis

Figure 3A shows that the model was most sensitive to *k_b_* for *T*_1,*b*_ = 0.8 s and PLD = 1.5 s. These optimal values provided over a 3-fold increase in sensitivity compared to the use of no contrast (equivalent to *T*_1,*b*_ = 1.65 s). Sensitivity remained within 90% of the maximum value over the ranges 0.6 s ≤ *T*_1,*b*_ ≤ 1.0 s and 1.3 s ≤ PLD ≤ 1.9 s. Minimal sensitivity (under 10% of the maximum value) was observed for both very short *T*_1,*b*_ values (*T*_1,*b*_ ≤ 0.3 s) and for *T*_1,*b*_ ≈ *T*_1,*e*_.

**Figure 3.**
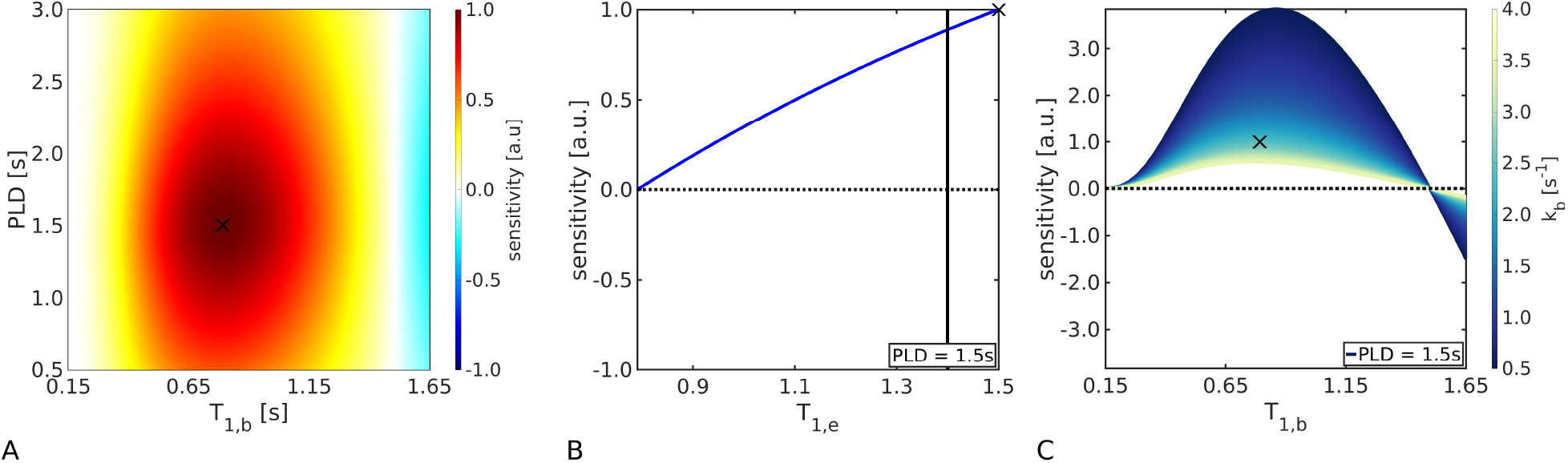
Sensitivity analyses. **(A)**. Sensitivity of the ASL difference signal to *k_b_* as a function of *T*_1,*b*_ and PLD (with *T*_1,*e*_ = 1.5 s, *k_b_* = 2.65 s^−1^). The colourbar shows the magnitude of the sensitivity functions, which have been normalised using the maximum value obtained over the range of parameter combinations (indicated by the black cross). **(B)**. Sensitivity dependence on *T*_1,*e*_, simulating the effect of extravasated contrast agent (with *T*_1,*b*_ = 0.8 s, PLD = 1.5 s, *k_b_* = 2.65 s^−1^). The black line indicates the potential *T*_1,*e*_ after leakage from minor BBB damage in vivo. **(C)**. Sensitivity dependence on underlying *k_b_* values (with *T*_1,*b*_ = 0.8 s, *T*_1,*e*_ = 1.5 s, PLD = 1.5 s). All sensitivity functions were normalised to the parameter set consisting of *k_b_* = 2.65 s^−1^, *T*_1,*b*_ = 0.8 s, *T*_1,*e*_ = 1.5 s and PLD = 1.5 s (indicated in each figure by the black cross). Other fixed parameters were: *f* = 60 ml blood / min / 100 ml tissue, *t_L_* = 2 s, *t_A_* = 1.2 s, λ = 0.9 and *α* = 0.85.

Reductions in *T*_1,*e*_ arising from extravasated contrast agent corresponded to an approximately linear decrease in sensitivity, culminating in zero sensitivity to *k_b_* for *T*_1,*e*_ = *T*_1,*b*_ (Figure 3B). This represents the full range of BBB integrity, from fully intact with no leakage of the GBCA (i.e. *T*_1,*e*_ = 1.5 s, the equilibrium value) to unobstructed leakage (i.e. *T*_1,*e*_ = *T*_1,*b*_). A reduction of ~0.1 s may be expected for minor BBB damage (34), corresponding to a decrease in sensitivity of ~ 10 %.

Figure 3C shows the sensitivity dependence of the model to underlying *k_b_* values. Greater sensitivity was observed for slower exchange rates. From the magnitude of the sensitivity function - which provides an indication of the expected level of measurement precision for a given noise level - it can be seen that, compared to a baseline *k_b_* = 2.65 s^−1^, an increase of 15% in the exchange rate to *k_b_* = 2.92 s^−1^ would correspond to a 12% reduction in measurement precision. The optimal *T*_1,*b*_ varied minimally from *T*_1,*b*_ = 0.77 s at the fastest exchange rate (*k_b_* = 4.0 s^−1^) to *T*_1,*b*_ = 0.86 s at the slowest exchange rate (*k_b_* = 0.5 s^−1^).

### Error propagation

Figure 4 shows the errors propagated into *k_b_*, *f* and *t_A_* by errors in *T*_1_ values. The accuracy of *k_b_* was highly sensitive to errors in both 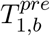 and 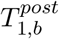: to obtain estimates of *k_b_* with less than 10% error required 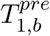 to be known within ±1. 5% and 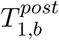 within ±0.7 %. Errors in *T_1_,_e_* propagated less uncertainty into *k_b_* estimates, requiring a measurement accuracy of ±11% to maintain the same 10% error level in *k_b_*.

**Figure 4.**
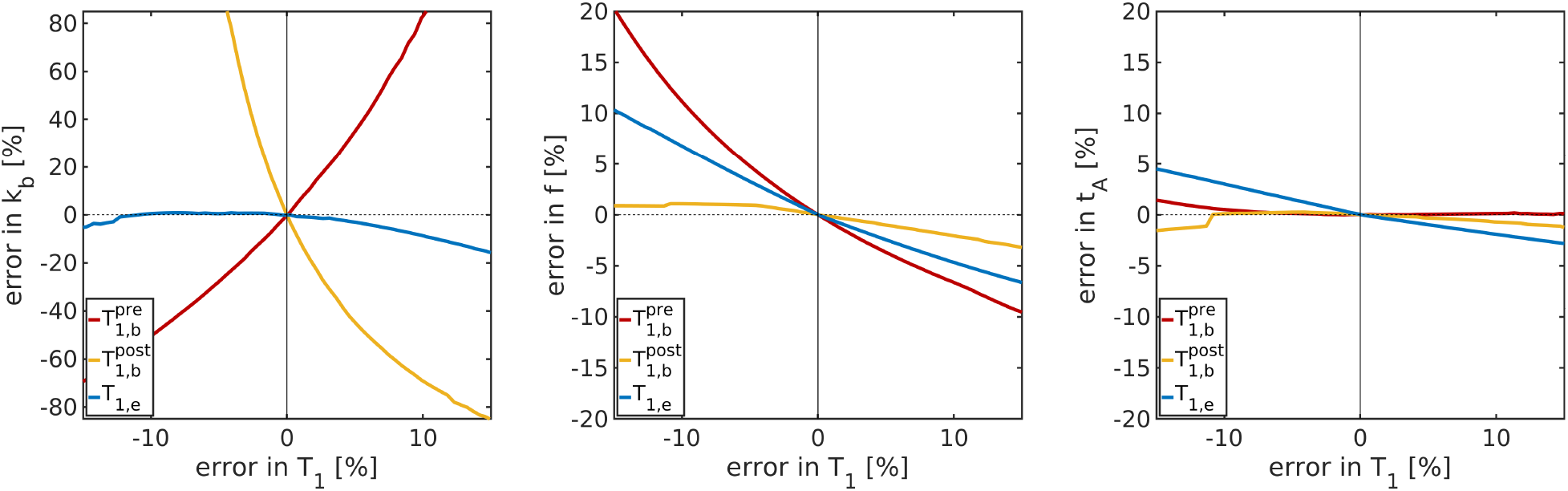
Error propagation. The error propagated into fitted parameters from errors in *T*_1_ values is shown for the exchange rate (*k_b_*, left), CBF (*f*, centre) and ATT (*t_A_*, right). Ground truth parameter values were: *k_b_* = 2.65 s^−1^, *f* = 60 ml blood / min / 100 ml tissue, *t_A_* = 1.2 s, 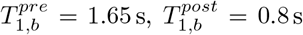, *T*_1,*e*_ = 1.5 s, 5 pre-contrast PLDs between 0.9 – 2.1 s, 1 post-contrast PLD = 1.5 s, *t_L_* = 2 s, λ = 0.9 and *α* = 0.85.

CBF accuracy was more influenced by errors in 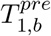 and *T*_1,*e*_ than in 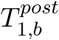. To estimate *f* with under 10% error required a measurement accuracy within ±9.1% for 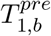 and within ±14.4% for *T*_1,*e*_. The CBF error was under 5% for all simulated 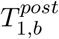 errors (±15 %).

The error propagated into ATT estimates was under 5% for all simulated *T*_1_ errors (±15 %).

### Monte Carlo simulations

The variation in accuracy and precision of fitted model parameters with underlying *k_b_* values is shown in Figure 5A. No biases were evident in any of the *k_b_* estimates; however, precision, indicated by the shaded error bars (IQR of fitted values), was reduced at higher *k_b_* values, as predicted by the sensitivity analysis in Figure 3C. The accuracy and precision of CBF and ATT were largely unaffected by underlying exchange rates. The number of extreme fits was under 6% in all cases (see Supporting Information, Figure S1).

**Figure 5.**
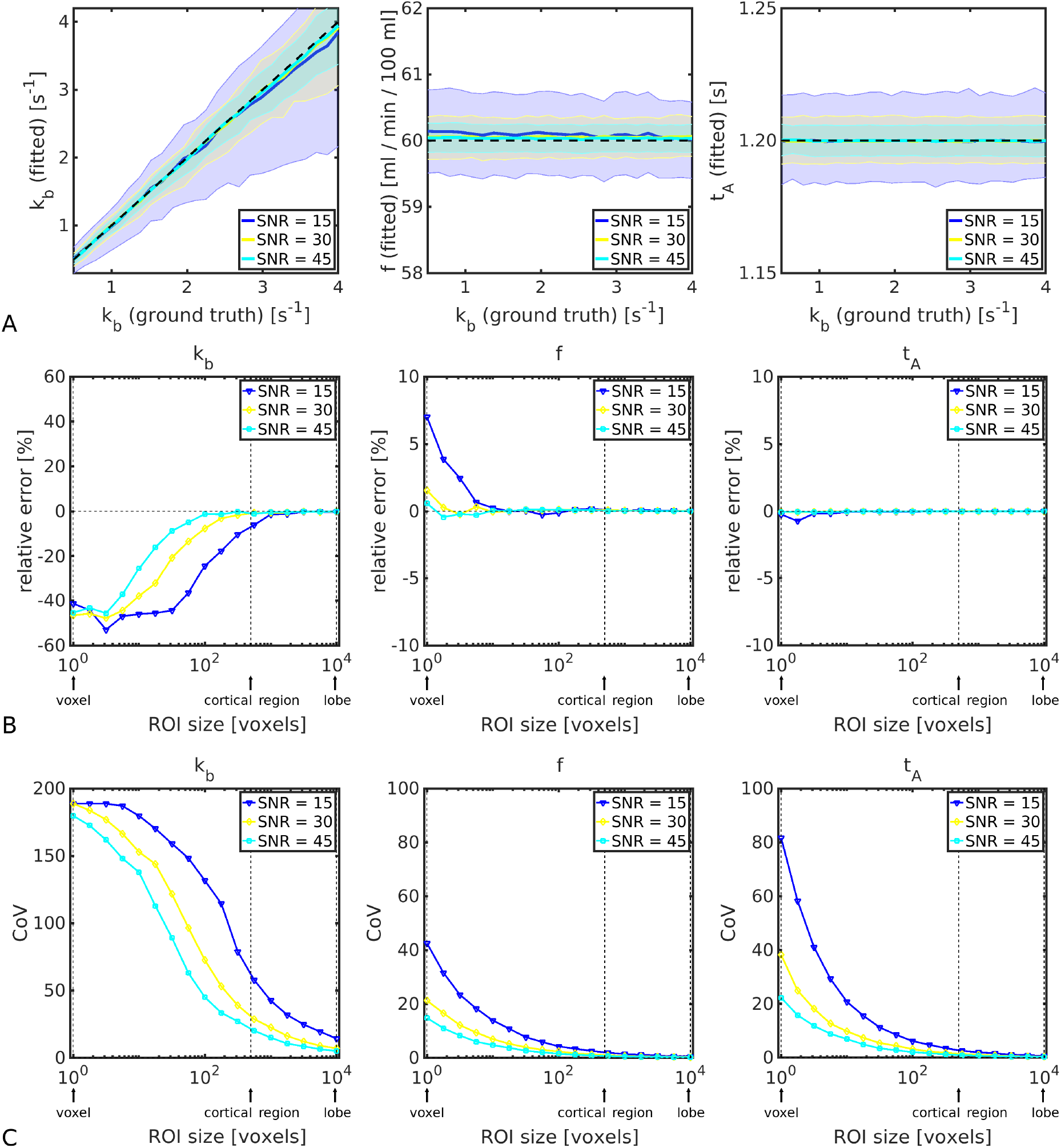
Monte Carlo simulations. **(A)**. Median parameter values (solid lines) for a simulated cortical ROI (500 voxels) are shown as a function of ground truth exchange rate for the fitted exchange rate (*k_b_*, left), CBF (*f*, centre), and ATT (*t_A_*, right). Shaded regions indicate the IQR of fitted values; black dashed lines indicate ground truth parameter values. **(B)**. Relative errors in parameter estimates after signal averaging across different simulated ROI sizes are shown for the exchange rate (*k_b_*, left), CBF (*f*, centre), and ATT (*t_A_*, right). **(C)**. The CoV of fitted parameters are shown for the exchange rate (*k_b_*, left), CBF (f, centre), and ATT (*t_A_*, right). Displayed SNR levels indicate voxel-wise values in the control signal. Full model parameter details are provided in Table 2.

The feasibility of regional water exchange measurements is considered in Figure 5B-C. Given a voxel-wise SNR of 30 in the control signal (and fixed *k_b_* = 2.65 s^−1^), in a cortical ROI (500 voxels) the relative error of *k_b_* was under 1% and the CoV was 30 %. Signal averaging across a simulated lobe (10000 voxels) reduced the CoV to 7 %. The CoV of voxel-level *k_b_* estimates was very high (190 %). CBF and ATT were estimated with good accuracy (relative error *ϵ* < 1 %) and reasonable precision (COV_*f*_ < 21% and COV*t_A_* < 38 %) at the voxel level for SNR = 30.

### Optimal injected GBCA dose

Figure 6 shows the recovery of 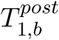 with time after contrast agent injection. Optimal values of 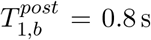 were obtained approximately 3 min after injection of a 0.025 mmol/kg dose (quarter dose), 21 min after a 0.050 mmol/kg dose (half dose), 37 min after a 0.075 mmol/kg dose (three quarter dose) and 48 min after a 0.100 mmol/kg dose (full dose).

**Figure 6.**
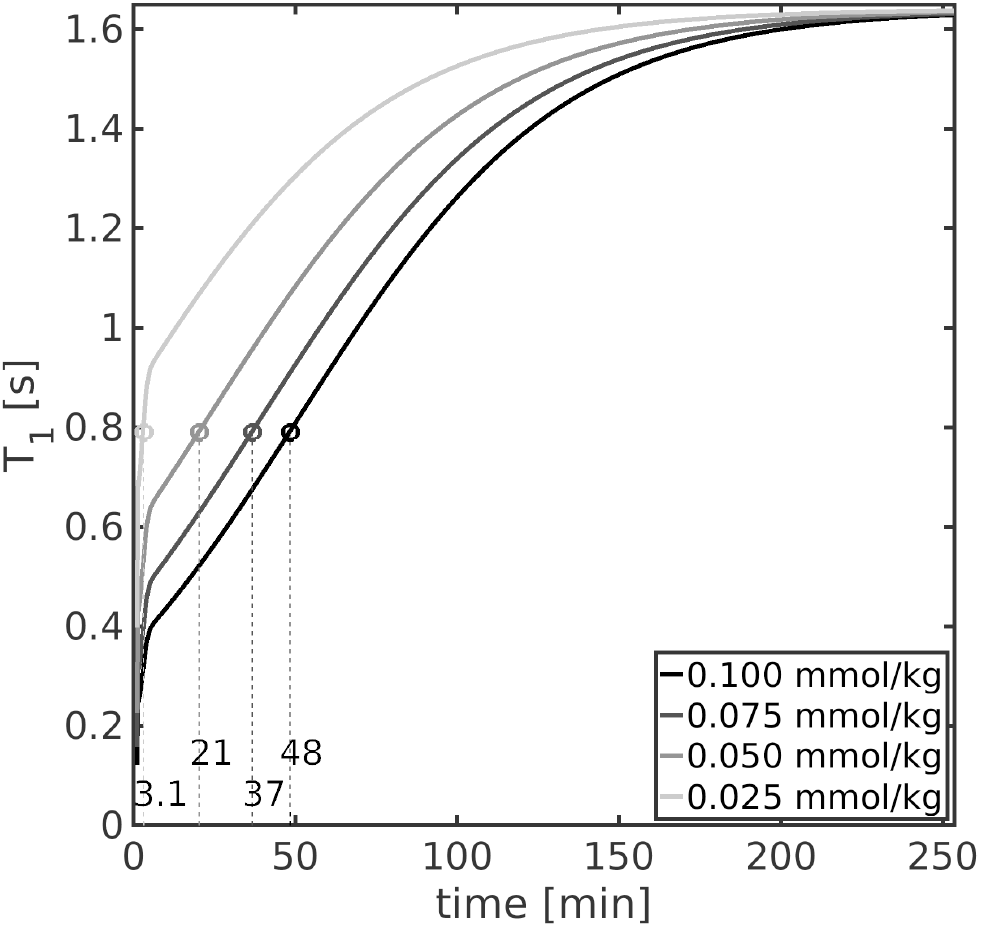
Optimal injected GBCA dose. Blood *T*_1_ recovery curves post-contrast are shown for different dose levels; a standard full dose is 0.100 mmol/kg. The dotted lines indicate the time post injection to reach the optimal 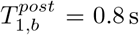, highlighted for each dose level by the circles.

### In vivo data

The mean and SD across subjects of GM *T*_1_ values were: (i) *T*_1,*e*_ = 1.50 ± 0.09 s pre-contrast; (ii) *T*_1,*e*_ = 1.48 ± 0.09 s for PC1, and; (iii) *T*_1,*e*_ = 1.39 ± 0.09 s for PC2. Representative tissue *T*_1_ maps pre- and post-contrast are provided in the Supporting Information, Figure S3. Pre- and post-contrast blood *T*_1,*b*_ values were: (i) 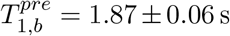 and 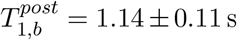 for PC1, and; (ii) 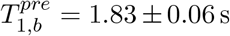 and 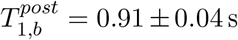 for PC2. As the PC2 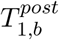 best approximated the optimal value, results from the PC2 data set will primarily be presented from here on; results from the PC1 data can be found in the Supporting Information (Figure S6.1 and Table S6.2).

Figure 7 shows parameter maps derived from the PC2 data for all 6 subjects. Mean regional parameter estimates for a selection of 18 cortical and sub-cortical regions (of particular relevance to dementia) are provided in Figure 8; results from all 90 regions can be found in the Supporting Information, Table S2. Good left/right hemispheric correspondence was observed in the exchange rate maps, although a few extreme fits (*k_b_* < 0 s^−1^ or *k_b_* > 10 s^−1^) were noted in some subjects. Averaged across subjects, the mean and SD of parameter values were: *t_A_* = 1.15 ± 0.49 s, *f* = 58.0 ± 14.3 ml blood / min / 100 ml tissue, *k_b_* = 2.32 ± 2.49 s^−1^.

**Figure 7.**
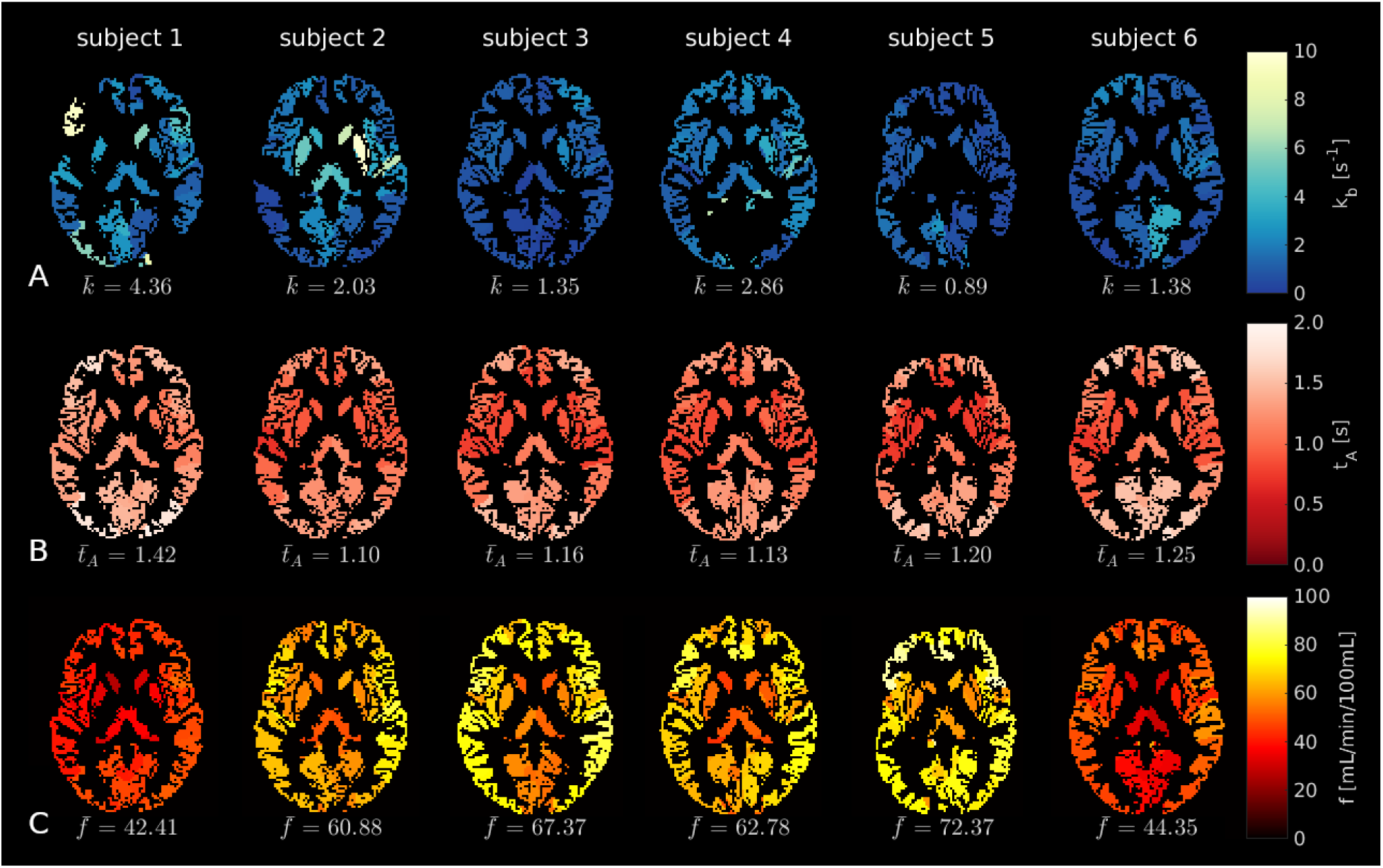
ASL parameter maps (PC2). **(A)**. Exchange rate, *k_b_*. **(B)**. Arterial transit time, *t_A_*. **(C)**. Cerebral blood flow, *f*. In all maps, black voxels represent masked white matter and CSF, as well as extreme *k_b_* fits (i.e. *k_b_* < 0 s^−1^ or *k_b_* > 10 s^−1^).

**Figure 8.**
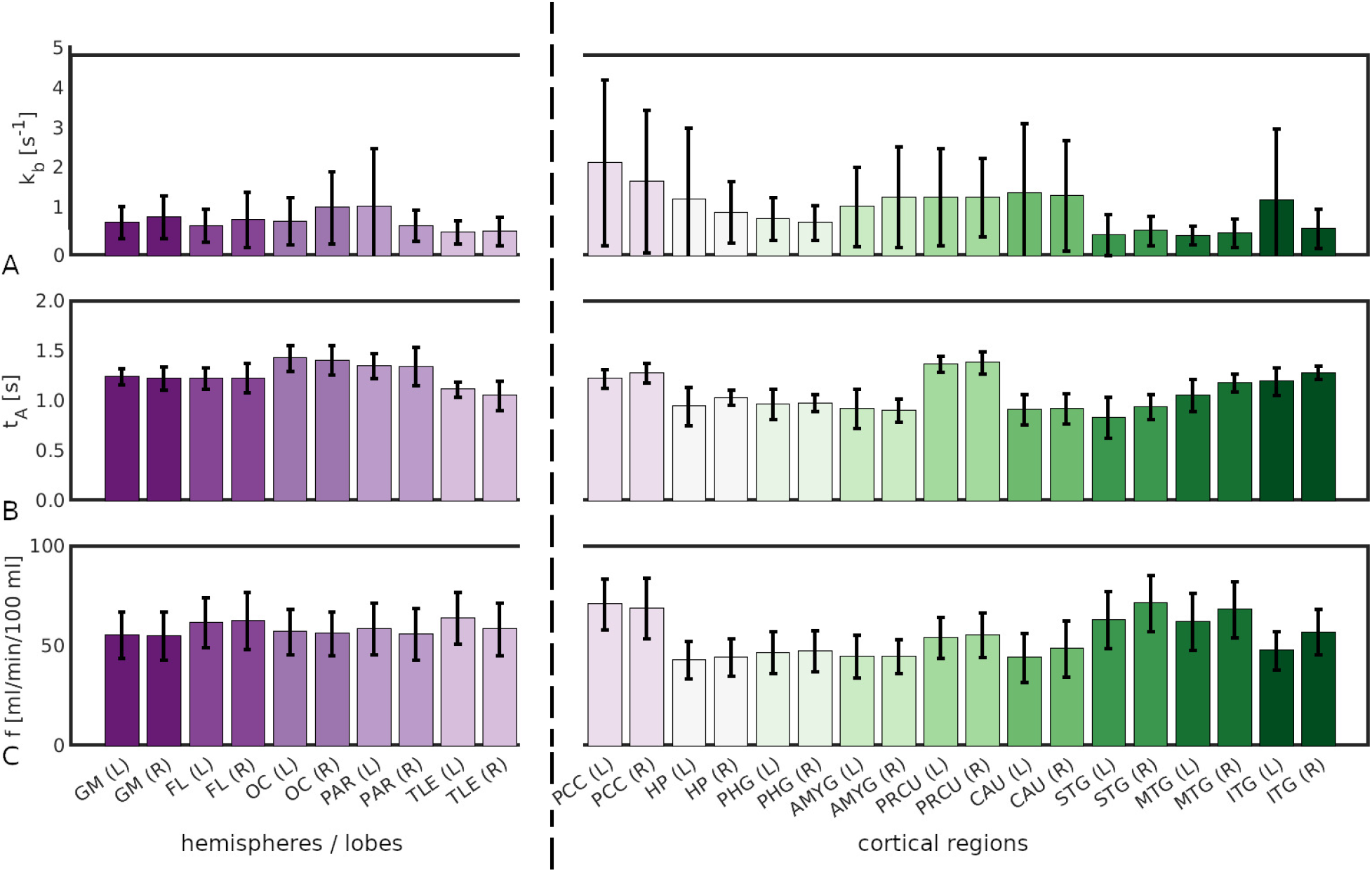
Mean parameter values across subjects in selected regions. **(A)**. Exchange rate, *k_b_*. **(B)**. Arterial transit time, *t_A_*; **(C)**. Cerebral blood flow, *f*. In all subplots, bar height represents the mean and error bars show the standard deviation across subjects.

Example ASL subtraction images pre- and post-contrast are shown in the Supporting Information, Figure S3. The mean SNR across segmented regions in the subtraction image at PLD = 1.5 s was 3.6 pre-contrast and 3.1 in the PC2 data (corresponding to SNR ~ 30 in the unlabelled data in simulations), indicating that the increased number of averages post-contrast compensated well for the expected loss of signal.

## Discussion

This simulation and proof-of-concept study demonstrates that measurements of BBB permeability to water are feasible using CE-ASL if accurate *T*_1_ values can be obtained: under the influence of an intravascular GBCA, the increased difference between blood water and tissue *T*_1_ relaxation times enables the signal contribution from intra- and extravascular compartments to be distinguished and *k_b_* to be estimated.

Identifying the optimal difference between blood water and tissue *T*_1_ relaxation times is key to obtaining reliable water exchange estimates using this method owing to the inherent trade-off between the reduction in 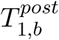 and sensitivity to *k_b_* (Figure 3C). Marginal reductions in 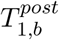 do not sufficiently perturb the post-contrast signal, meaning that the relative contributions of intra- and extravascular compartments remain difficult to separate and sensitivity is correspondingly low. Conversely, extreme reductions in 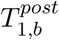 lead to a vanishing difference signal as fast recovery of the labelled spins relative to the ATT negates the effect of the inversion, rendering the post-contrast signal equivalent to the control data. It was shown using simulations that moderate reductions in 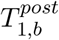 best enabled water exchange measurements, and, moreover, that a range of 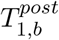 presented a similar capacity to reliably estimate *k_b_*. Practically this allows for flexibility in protocol design, as precise timings of the post-contrast ASL acquisition relative to GBCA administration are not necessary; however, increasing the number of signal averages for lower 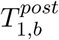 values may be prudent as the difference signal will be smaller and more susceptible to noise. Simulations showed that even a quarter dose (0.025 mmol/kg) could provide the optimal 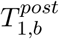 (Figure 6), which has reduced safety concerns compared to a full dose.

The sensitivity analysis indicated that more precise *k_b_* estimates can be expected at lower exchange rates (Figure 3C), meaning that CE-ASL is best suited to probing early, subtle damage. Monte Carlo simulations supported this finding and quantified the measurement precision of model parameters in terms of ground truth *k_b_* (Figure 5A). The accuracy and precision of fitted parameters was also quantified as a function of SNR (Figure 5B-C). CBF and ATT were estimated at the voxel level with acceptable precision at realistic noise levels in synthetic data. Reliable estimates of *k_b_* at the voxel level proved unfeasible, with a measurement precision approaching CoV ~ 190% (Figure 5C): inherently low SNR data combined with low sensitivity of the model to *k_b_* (relative to the CBF and ATT) renders *k_b_* a challenging parameter to fit. ROI analyses are valuable in this instance as the SNR effectively increases as the square root of the number of voxels, reducing random measurement errors and making regional *k_b_* estimates possible (Figure 7A). Practically, it should be noted that an upper limit on the ROI size used for regional analysis is likely to exist owing to variability in ATT, CBF and potentially *k_b_* across the brain, particularly in disease.

Regional in vivo *k_b_* measurements (Figure 8) were in agreement with literature values. The mean value across all segmented ROIs and subjects was *k_b_* = 2.32 s^−1^; previous studies have reported average GM values in the range 0.63 – 3.68 s^−1^ (13). There was also good agreement in regional values between hemispheres, suggesting that physiologically plausible *k_b_* values can be obtained using CE-ASL. CBF was notably lower in two subjects, although in line with published values (29; 41; 42) and reports of high inter-subject variability (43). It is also possible that age and gender were a factor: one of the subjects was the only male and one was older than the rest, and lower CBF has been reported in both demographics (44).

There are two limiting factors in the Monte Carlo simulations in this study. Firstly, regional variations in ATT observed in vivo were not modelled in the simulations: regions with *t_A_* > 1.5 s (i.e. ATT longer than the post-contrast PLD) are likely to have lower precision as the complete bolus may not have arrived at the voxel at the time of imaging. Acquiring data at multiple PLDs post-contrast may further ensure adequate sensitivity for regions with longer ATT, although a single post-contrast PLD = 1.5 s was chosen here to allow time for multiple signal averages. Secondly, alterations in *T*_1,*e*_ were not modelled in the simulations: contrast agent leakage into the extravascular space acts to decrease the difference between the post-contrast blood water and tissue *T*_1_ times, making the separation of intra- and extravascular signal components more challenging. The sensitivity analysis confirmed this, showing that, as 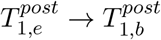, *k_b_* was estimated with increasingly poorer precision. In our proof-of-concept study a reduction of ~ 7% in tissue *T*_1_ was observed; however, this is the total tissue value and includes the blood component, which likely explains the majority of this decrease. Moreover, the decrease was consistent across subjects and so does not suggest leakage due to pathology. This raises the more general point that *T*_1,*e*_ is not independent of *k_b_*; however, this was also not modelled in simulations and should be considered in future studies. In the in vivo data, adding vascular crushers to the acquisition may improve estimation of the ATT and CBF: without vascular crushers, the signal at short PLDs may contain contributions from large vessels, leading to shorter ATT and higher CBF; however, as no exchange is expected to occur in large vessels, *k_b_* is unlikely to be affected.

The primary limitation of CE-ASL is the accuracy required in *T*_1_ measurements (Figure 4). Similar systematic errors in both 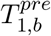 and 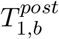 may mitigate error propagation into *k_b_* to some extent as opposing effects are introduced (see Supporting Information, Figure S4); however, particularly for 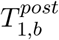, where sensitivity to *k_b_* and therefore error propagation is greatest, small errors can introduce significant biases into *k_b_* measurements. The potential effect on in vivo parameter estimates arising from *T*_1,*b*_ biases was explored post hoc by perturbing the measured 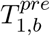 and 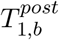 by ±10% and re-fitting the model (see Supporting Information, Figure S5). Mean fitted parameter values varied according to the trends predicted in Figure S4, with *k_b_* increased on average by 100% for *T*_1,*b*_ adjusted 10% lower and reduced by 44% for *T*_1,*b*_ adjusted 10% higher than the measured value. It must also be considered that inter-subject variability in *T*_1,*b*_ can be introduced depending on haematocrit levels and oxygen extraction fraction (45), which further emphasises the need for reliable individual *T*_1,*b*_ mapping; however, regional *k*b** variations within a subject may still be identified. Finally, clinical conditions in which key assumptions of the model are violated - for example arteriovenous malformations, where the passage of blood into the microvasculature is disrupted and the assumption of no outflow no longer holds - must be treated cautiously.

Given current clinical capabilities, these *T*_1_ accuracy requirements limit the utility of the CE-ASL technique at this point. Assuming a situation where *T*_1_ values are accurate enough, a clinically practical application of CE-ASL would be in conjunction with conventional DCE-MRI studies: ASL data acquired before and after a DCE-MRI protocol could utilise the residual effects of the GBCA to obtain the optimally-shortened 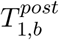 needed for CE-ASL imaging, and dose calculations suggest that the time to optimal 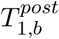 following a full-dose injection could make this approach feasible (32 min to reach the lower bound 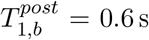). In cases where BBB damage is minor and DCE-MRI does not show significant uptake of the contrast agent in the tissue (3), concomitant acquisition of CE-ASL data could provide a complementary indication of subtle BBB breakdown. However, it is necessary to reduce the dependence of *k_b_* estimates on measured *T*_1_ values for this to be clinically viable.

## Abbreviations

ASL: arterial spin labelling
ATT: arterial transit time
BBB: blood-brain barrier
CASL: continuous arterial spin labelling
CBF: cerebral blood flow
CE-ASL: contrast-enhanced arterial spin labelling
DCE: dynamic contrast-enhanced
GBCA: gadolinium-based contrast agent
LD: labelling duration
PLD: post-labelling delay

## Acknowledgments

This work was supported by EPSRC grants EP/S031510/1 and EP/M005909/1. Thanks to GE Healthcare for use of the Enhanced ASL sequence.

## Appendices

## Appendix A: sensitivity functions

The sensitivity functions are defined as the partial derivative of the signal model with respect to *k_b_*; the shape and magnitude of the functions therefore indicate the sensitivity of the signal model to changes in *k_b_*. The sensitivity functions for the CASL model in Equation 4 are:

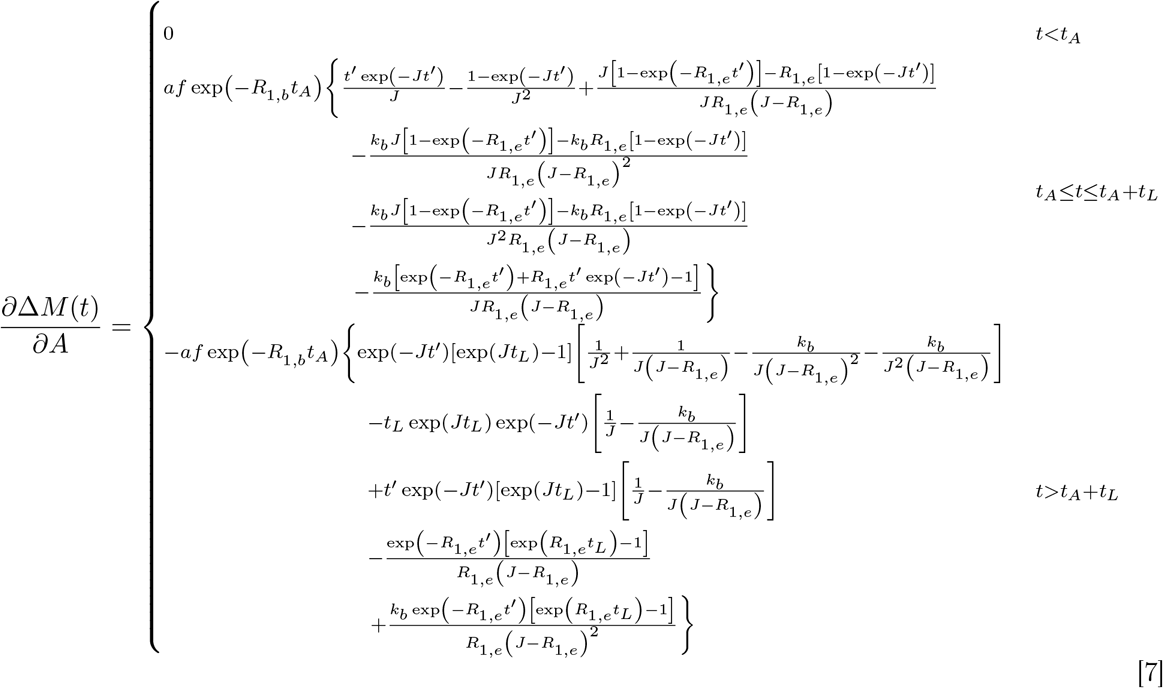

where *t_L_* is the labelling time, *t_A_* is the arterial transit time, *f* is the cerebral blood flow, *α* is the inversion efficiency of the labelling, *R*_1,*e*_ = ^1^/_*T*_1,*e*__, *R*_1,*b*_ = ^1^/_*T*_1,*b*__, *J* = *k_b_* + *R*_1,*b*_ and *t*′ = *t* – *t_A_*.

## Notes

### Competing Interest Statement

The authors have declared no competing interest.

