## Supporting information for "Blood-brain barrier water exchange measurements using contrast-enhanced ASL"

### Supporting information S1: extreme fits (synthetic data)

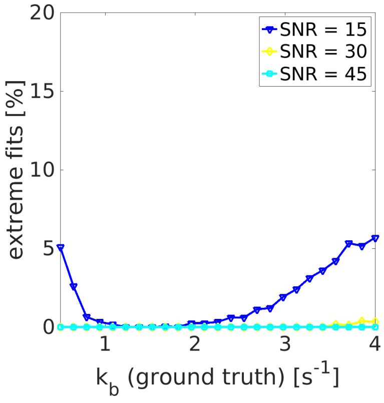


**Figure S1. Number of extreme fits (synthetic data).**

The number of extreme fits (defined as any parameter having hit a bound) in Monte Carlo simulations. Ground truth parameter values were: $k_b = 2.65\,\mathrm{s}^{-1}$, $f = 60\,\mathrm{ml\,blood} / \mathrm{min} / 100\,\mathrm{ml\,tissue}$, $t_A = 1.2\,\mathrm{s}$, $T_{1,b}^{pre} = 1.65\,\mathrm{s}$, $T_{1,b}^{post} = 0.8\,\mathrm{s}$, $T_{1,e} = 1.5\,\mathrm{s}$, 5 pre-contrast PLDs between $0.9 - 2.1\,\mathrm{s}$, 1 post-contrast PLD $= 1.5\,\mathrm{s}$, $t_L = 2\,\mathrm{s}$, $\lambda = 0.9$ and $\alpha = 0.85$.

### Supporting information S2: regional ASL parameter fits (PC2)

**Table S2. ASL regional parameter fits (PC2).**

Values are the mean and standard deviation across subjects of regional fits. GM = grey matter; FL = frontal lobe; OC = occipital lobe; PAR = parietal lobe; TLE = temporal lobe; PrcG = precentral gyrus; SFG = superior frontal gyrus; OFGsup = superior orbitofrontal gyrus; MFG = middle frontal gyrus; OFGmid = middle orbitofrontal gyrus; IFGop = inferior frontal gyrus (opercular part); IFGtr = inferior frontal gyrus (triangular part); OFGinf = inferior orbitofrontal gyrus; ROL = rolandic operculum; SMA = supplementary motor area; OLF = olfactory cortex; SFGmed = superior frontal gyrus (medial part); OFGmid = orbitofrontal gyrus (middle part); REC = rectus; INS = insula; ACC = anterior cingulate cortex; MCC = middle cingulate cortex; PCC = posterior cingulate cortex; HP = hippocampus; PHG = parahippocampal gyrus; AMYG = amygdala; CAL = calcarine; CUN = cuneus; LING = lingual gyrus; SOG = superior occipital gyrus; MOG = middle occipital gyrus; IOG = inferior occipital gyrus; FFG = fusiform gyrus; PocG = postcentral gyrus; SPG = superior parietal gyrus; IPL = inferior parietal lobule; SMG = supramarginal gyrus; ANG = angular gyrus; PRCU = precuneus; PCL = paracentral lobule; CAU = caudate; PUT = putamen; PAL = pallidum; THAL = thalamus; HES = Heschl gyrus; STG = superior temporal gyrus; TPsup = temporal pole (superior part); MTG = middle temporal gyrus; TPmid = temporal pole (middle part); ITG = inferior temporal gyrus; L = left; R = right.

| Region name | No. voxels | $f$ [ml / min / 100 ml] | $t_A$ [s] | $k_b$ [$s^{-1}$] |
|---|---|---|---|---|
| GM (L) | 55805 | $55.53 \pm 12.30$ | $1.25 \pm 0.09$ | $1.67 \pm 0.84$ |
| GM (R) | 46429 | $55.25 \pm 12.75$ | $1.23 \pm 0.12$ | $1.93 \pm 1.06$ |
| FL (L) | 12154 | $61.88 \pm 12.73$ | $1.23 \pm 0.12$ | $1.51 \pm 0.91$ |
| FL (R) | 11744 | $62.78 \pm 15.11$ | $1.23 \pm 0.15$ | $1.81 \pm 1.47$ |
| OC (L) | 6678 | $57.16 \pm 12.67$ | $1.43 \pm 0.15$ | $1.74 \pm 1.12$ |
| OC (R) | 7591 | $56.36 \pm 12.16$ | $1.41 \pm 0.16$ | $2.42 \pm 1.72$ |
| PAR (L) | 4287 | $58.56 \pm 12.59$ | $1.35 \pm 0.14$ | $2.49 \pm 3.19$ |
| PAR (R) | 4561 | $55.96 \pm 12.75$ | $1.35 \pm 0.21$ | $1.52 \pm 0.82$ |
| TLE (L) | 7046 | $64.23 \pm 12.79$ | $1.12 \pm 0.08$ | $1.18 \pm 0.61$ |
| TLE (R) | 6643 | $58.61 \pm 12.69$ | $1.05 \pm 0.17$ | $1.24 \pm 0.77$ |
| PrcG (L) | 1379 | $58.74 \pm 16.53$ | $1.37 \pm 0.33$ | $2.86 \pm 3.63$ |
| PrcG (R) | 1212 | $57.32 \pm 12.20$ | $1.35 \pm 0.24$ | $1.40 \pm 1.09$ |
| SFG (L) | 1366 | $56.18 \pm 14.20$ | $1.39 \pm 0.30$ | $1.56 \pm 0.74$ |
| SFG (R) | 1696 | $57.86 \pm 13.22$ | $1.40 \pm 0.16$ | $1.34 \pm 0.84$ |
| OFGsup (L) | 494 | $56.60 \pm 16.07$ | $1.20 \pm 0.09$ | $1.95 \pm 1.33$ |
| OFGsup (R) | 511 | $53.32 \pm 7.75$ | $1.22 \pm 0.12$ | $2.73 \pm 2.87$ |

| Region name | No. voxels | $f$ | $t_A$ | $k_b$ |
|---|---|---|---|---|
| MFG (L) | 2142 | $64.83 \pm 14.75$ | $1.45 \pm 0.32$ | $2.82 \pm 3.61$ |
| MFG (R) | 2293 | $65.59 \pm 14.69$ | $1.42 \pm 0.12$ | $1.60 \pm 0.81$ |
| OFGmid (L) | 428 | $64.46 \pm 14.86$ | $1.32 \pm 0.11$ | $2.52 \pm 1.77$ |
| OFGmid (R) | 471 | $62.80 \pm 14.10$ | $1.28 \pm 0.14$ | $4.79 \pm 3.48$ |
| IFGop (L) | 439 | $72.95 \pm 16.65$ | $1.08 \pm 0.15$ | $2.88 \pm 3.54$ |
| IFGop (R) | 615 | $73.92 \pm 14.68$ | $1212 \pm 0.12$ | $1.86 \pm 1.41$ |
| IFGtr (L) | 1024 | $69.88 \pm 17.41$ | $1.18 \pm 0.17$ | $2.91 \pm 3.20$ |
| IFGtr (R) | 808 | $68.27 \pm 13.45$ | $1100 \pm 0.14$ | $1.84 \pm 1.53$ |
| OFGinf (L) | 856 | $68.32 \pm 17.04$ | $1.08 \pm 0.20$ | $2.70 \pm 3.63$ |
| OFGinf (R) | 768 | $62.89 \pm 11.04$ | $0.93 \pm 0.27$ | $1.82 \pm 1.20$ |
| ROL (L) | 514 | $50.75 \pm 13.04$ | $0.99 \pm 0.22$ | $2.96 \pm 3.46$ |
| ROL (R) | 689 | $58.11 \pm 12.05$ | $1.00 \pm 0.15$ | $1.80 \pm 1.22$ |
| SMA (L) | 905 | $58.35 \pm 15.36$ | $1.18 \pm 0.15$ | $1.67 \pm 1.42$ |
| SMA (R) | 1016 | $54.88 \pm 13.61$ | $1.19 \pm 0.18$ | $1.14 \pm 0.69$ |
| OLF (L) | 187 | $58.25 \pm 14.52$ | $0.87 \pm 0.11$ | $1.79 \pm 1.03$ |
| OLF (R) | 189 | $55.75 \pm 14.21$ | $0.91 \pm 0.16$ | $2.06 \pm 1.31$ |
| SFGmed (L) | 1162 | $63.80 \pm 15.38$ | $1.08 \pm 0.14$ | $1.24 \pm 1.02$ |
| SFGmed (R) | 983 | $63.10 \pm 13.73$ | $1.15 \pm 0.13$ | $1.22 \pm 0.90$ |
| OFGmed (L) | 370 | $74.18 \pm 17.43$ | $1.00 \pm 0.17$ | $1.10 \pm 0.49$ |
| OFGmed (R) | 463 | $71.02 \pm 15.17$ | $1.02 \pm 0.16$ | $1.86 \pm 1.58$ |
| REC (L) | 478 | $70.97 \pm 17.53$ | $0.93 \pm 0.15$ | $0.81 \pm 0.53$ |
| REC (R) | 440 | $62.54 \pm 11.04$ | $0.99 \pm 0.12$ | $1.89 \pm 1.86$ |
| INS (L) | 1243 | $57.39 \pm 12.48$ | $0.92 \pm 0.21$ | $2.91 \pm 3.83$ |
| INS (R) | 1097 | $61.62 \pm 12.25$ | $0.85 \pm 0.15$ | $1.80 \pm 1.11$ |
| ACC (L) | 812 | $71.08 \pm 16.71$ | $0.90 \pm 0.14$ | $0.99 \pm 0.24$ |
| ACC (R) | 753 | $67.65 \pm 18.12$ | $0.96 \pm 0.17$ | $2.82 \pm 3.83$ |
| MCC (L) | 1095 | $62.56 \pm 14.11$ | $1.10 \pm 0.10$ | $2.09 \pm 0.99$ |
| MCC (R) | 1292 | $63.89 \pm 15.44$ | $1.09 \pm 0.18$ | $2.44 \pm 1.57$ |
| PCC (L) | 192 | $71.00 \pm 14.32$ | $1.23 \pm 0.09$ | $4.66 \pm 3.91$ |
| PCC (R) | 110 | $69.02 \pm 16.79$ | $1.28 \pm 0.11$ | $3.72 \pm 3.60$ |
| HP (L) | 614 | $43.05 \pm 9.19$ | $0.95 \pm 0.21$ | $2.85 \pm 3.82$ |
| HP (R) | 530 | $44.37 \pm 10.10$ | $1.03 \pm 0.08$ | $2.18 \pm 1.72$ |
| PHG (L) | 555 | $46.72 \pm 11.57$ | $0.97 \pm 0.17$ | $1.85 \pm 1.04$ |
| PHG (R) | 698 | $47.49 \pm 11.41$ | $0.98 \pm 0.09$ | $1.66 \pm 0.96$ |
| AMYG (L) | 174 | $44.85 \pm 11.66$ | $0.92 \pm 0.22$ | $2.46 \pm 2.14$ |
| AMYG (R) | 187 | $44.83 \pm 9.29$ | $0.91 \pm 0.13$ | $2.93 \pm 2.68$ |
| CAL (L) | 1266 | $57.84 \pm 13.07$ | $1.36 \pm 0.14$ | $4.50 \pm 4.28$ |

| Region name | No. voxels | $f$ | $t_A$ | $k_b$ |
|---|---|---|---|---|
| CAL (R) | 1010 | $56.81 \pm 16.15$ | $1.35 \pm 0.13$ | $3.82 \pm 4.97$ |
| CUN (L) | 715 | $52.83 \pm 10.21$ | $1.42 \pm 0.13$ | $3.46 \pm 3.62$ |
| CUN (R) | 723 | $55.12 \pm 11.46$ | $1.48 \pm 0.19$ | $3.11 \pm 3.10$ |
| LING (L) | 1253 | $52.14 \pm 12.57$ | $1.28 \pm 0.12$ | $4.92 \pm 3.81$ |
| LING (R) | 1296 | $55.16 \pm 14.85$ | $1.31 \pm 0.12$ | $3.42 \pm 4.28$ |
| SOG (L) | 549 | $49.96 \pm 8.34$ | $1.59 \pm 0.20$ | $1.50 \pm 1.27$ |
| SOG (R) | 576 | $58.53 \pm 12.30$ | $1.69 \pm 0.28$ | $3.98 \pm 3.31$ |
| MOG (L) | 1716 | $62.60 \pm 11.65$ | $1.49 \pm 0.21$ | $1.81 \pm 2.15$ |
| MOG (R) | 1035 | $64.05 \pm 10.46$ | $1.54 \pm 0.16$ | $2.01 \pm 4.29$ |
| IOG (L) | 542 | $60.60 \pm 14.86$ | $1.47 \pm 0.25$ | $2.75 \pm 3.98$ |
| IOG (R) | 495 | $61.03 \pm 11.53$ | $1.50 \pm 0.18$ | $1.92 \pm 4.34$ |
| FFG (L) | 1562 | $41.84 \pm 10.54$ | $1.21 \pm 0.15$ | $3.54 \pm 3.42$ |
| FFG (R) | 1635 | $42.62 \pm 11.75$ | $1.24 \pm 0.06$ | $2.67 \pm 2.52$ |
| PoCG (L) | 1473 | $55.99 \pm 13.30$ | $1.35 \pm 0.24$ | $1.56 \pm 0.57$ |
| PoCG (R) | 1341 | $52.46 \pm 11.63$ | $1.34 \pm 0.14$ | $1.40 \pm 0.45$ |
| SPG (L) | 704 | $46.51 \pm 10.23$ | $1.60 \pm 0.21$ | $3.94 \pm 2.31$ |
| SPG (R) | 620 | $45.27 \pm 10.15$ | $1.61 \pm 0.22$ | $4.56 \pm 3.79$ |
| IPL (L) | 1165 | $55.17 \pm 12.85$ | $1.38 \pm 0.23$ | $2.68 \pm 3.90$ |
| IPL (R) | 583 | $59.78 \pm 12.99$ | $1.41 \pm 0.17$ | $2.82 \pm 3.96$ |
| SMG (L) | 599 | $61.77 \pm 15.25$ | $1.14 \pm 0.20$ | $1.08 \pm 0.32$ |
| SMG (R) | 921 | $67.24 \pm 14.54$ | $1.14 \pm 0.14$ | $2.83 \pm 3.91$ |
| ANG (L) | 621 | $62.50 \pm 14.26$ | $1.31 \pm 0.13$ | $1.27 \pm 0.74$ |
| ANG (R) | 822 | $66.29 \pm 13.12$ | $1.36 \pm 0.12$ | $2.06 \pm 1.93$ |
| PRCU (L) | 1551 | $54.17 \pm 11.16$ | $1.37 \pm 0.09$ | $2.94 \pm 2.36$ |
| PRCU (R) | 1544 | $55.52 \pm 12.24$ | $1.39 \pm 0.12$ | $2.94 \pm 1.74$ |
| PCL (L) | 420 | $44.50 \pm 7.98$ | $1.32 \pm 0.09$ | $1.38 \pm 1.76$ |
| PCL (R) | 287 | $46.85 \pm 10.76$ | $1.36 \pm 0.17$ | $1.07 \pm 0.94$ |
| CAU (L) | 545 | $44.22 \pm 13.49$ | $0.92 \pm 0.17$ | $3.13 \pm 3.77$ |
| CAU (R) | 584 | $48.77 \pm 15.66$ | $0.92 \pm 0.16$ | $3.01 \pm 2.86$ |
| PUT (L) | 674 | $51.64 \pm 10.90$ | $0.88 \pm 0.18$ | $2.49 \pm 1.45$ |
| PUT (R) | 690 | $50.68 \pm 9.07$ | $0.95 \pm 0.13$ | $3.53 \pm 3.44$ |
| PAL (L) | 71 | $52.74 \pm 11.07$ | $0.90 \pm 0.19$ | $3.62 \pm 2.56$ |
| PAL (R) | 67 | $54.81 \pm 12.35$ | $0.87 \pm 0.15$ | $1.98 \pm 1.30$ |
| THAL (L) | 357 | $48.35 \pm 11.15$ | $1.13 \pm 0.11$ | $1.54 \pm 1.88$ |
| THAL (R) | 399 | $46.75 \pm 11.94$ | $1.17 \pm 0.07$ | $1.64 \pm 1.72$ |
| HES (L) | 136 | $63.70 \pm 15.17$ | $-0.74 \pm 4.18$ | $2.19 \pm 4.26$ |
| HES (R) | 139 | $65.36 \pm 12.58$ | $0.87 \pm 0.11$ | $3.11 \pm 2.55$ |
| STG (L) | 1142 | $63.24 \pm 13.60$ | $0.83 \pm 0.23$ | $1.07 \pm 1.14$ |

| Region name | No. voxels | $f$ | $t_A$ | $k_b$ |
|---|---|---|---|---|
| STG (R) | 1474 | $71.43 \pm 13.68$ | $0.94 \pm 0.11$ | $1.26 \pm 0.76$ |
| TPsup (L) | 451 | $60.06 \pm 14.99$ | $0.85 \pm 0.15$ | $0.91 \pm 0.22$ |
| TPsup (R) | 515 | $60.52 \pm 14.11$ | $0.88 \pm 0.20$ | $1.33 \pm 1.13$ |
| MTG (L) | 2696 | $62.31 \pm 14.28$ | $1.06 \pm 0.18$ | $1.03 \pm 0.52$ |
| MTG (R) | 2427 | $68.43 \pm 14.11$ | $1.18 \pm 0.08$ | $1.13 \pm 0.80$ |
| TPmid (L) | 342 | $54.28 \pm 12.58$ | $1.06 \pm 0.09$ | $1.46 \pm 0.28$ |
| TPmid (R) | 482 | $51.26 \pm 8.23$ | $1.02 \pm 0.12$ | $1.44 \pm 0.75$ |
| ITG (L) | 1876 | $47.82 \pm 9.20$ | $1.20 \pm 0.15$ | $2.81 \pm 3.87$ |
| ITG (R) | 2009 | $57.09 \pm 11.50$ | $1.28 \pm 0.07$ | $1.37 \pm 1.02$ |

#### Supporting information S3: representative ASL subtraction images and tissue $T_1$ maps pre- and post-contrast

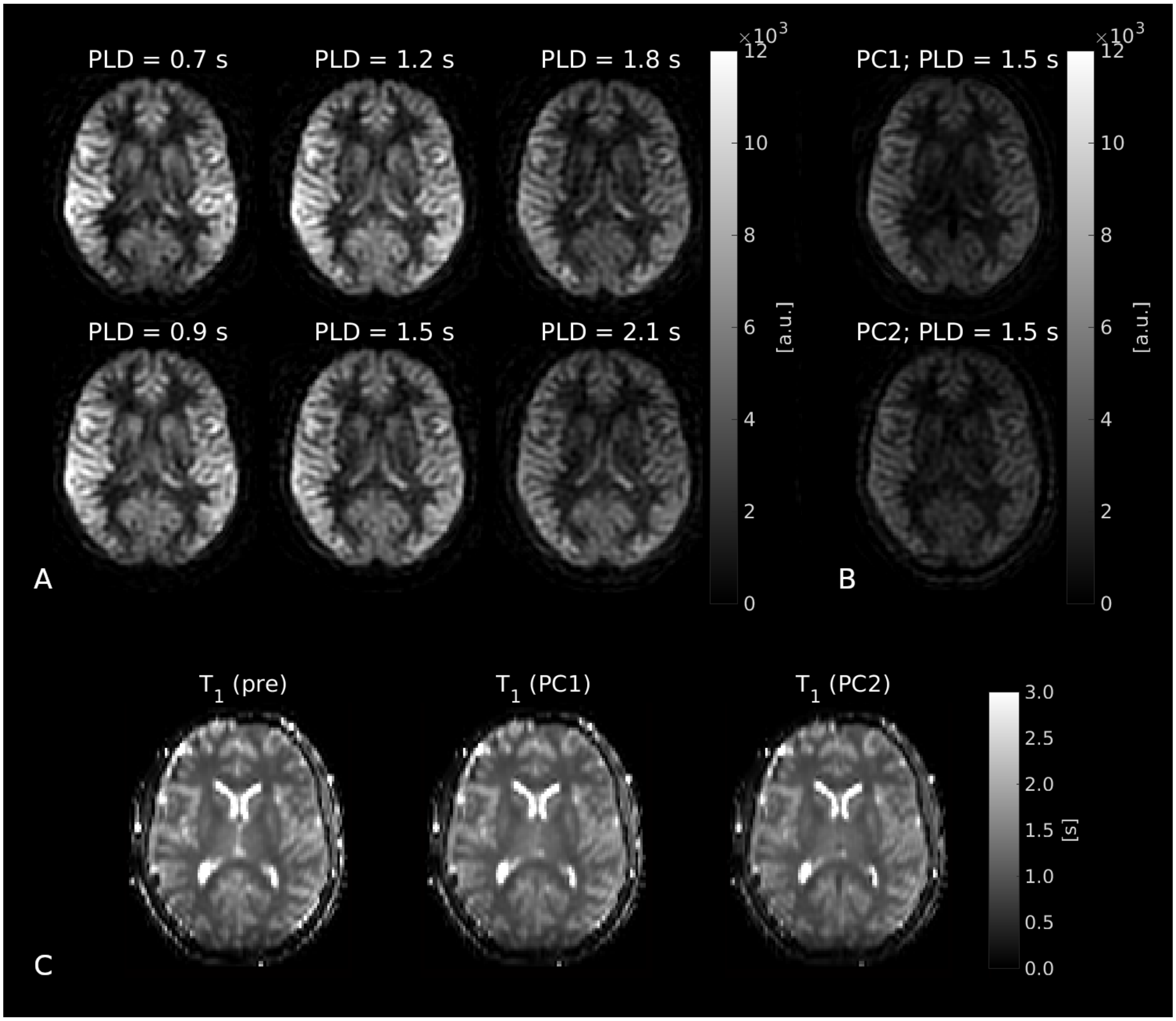


**Figure S3. Representative ASL data and tissue $T_1$ maps.**

**(A)**. ASL subtraction images pre-contrast at all post-labelling delay (PLD) times. The SNR at PLD = 1.5 s was 3.9. **(B)**. ASL subtraction data at the single PLD for the first (top; PC1) and second (bottom; PC2) post-contrast acquisitions. The SNR was 3.6 for PC1 and 3.1 for PC2. Good grey:white matter contrast remained visible following both contrast agent injections. **(C)**. Tissue $T_1$ maps pre- and post-contrast. Minimal difference was observed, indicating that the contrast agent remained in the intravascular space.

### Supporting information S4: systematic errors in blood $T_1$ values (synthetic data)

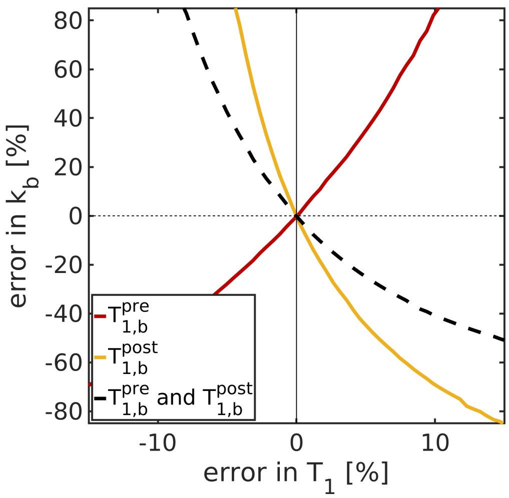


**Figure S4. Effect of systematic blood $T_1$ errors (synthetic data).**
The error propagated into $k_b$ estimates from systematic errors affecting $T_{1,b}^{pre}$ and $T_{1,b}^{post}$ equally (black dashed line); errors propagated into $k_b$ from errors in $T_{1,b}^{pre}$ and $T_{1,b}^{post}$ separately are shown again for completeness (red and yellow lines). Ground truth parameter values were: $k_b = 2.65\,\mathrm{s}^{-1}$, $f = 60\,\mathrm{ml\,blood}\,/\,\mathrm{min}\,/\,100\,\mathrm{ml\,tissue}$, $t_A = 1.2\,\mathrm{s}$, $T_{1,b}^{pre} = 1.65\,\mathrm{s}$, $T_{1,b}^{post} = 0.8\,\mathrm{s}$, $T_{1,e} = 1.5\,\mathrm{s}$, 5 pre-contrast PLDs between $0.9 - 2.1\,\mathrm{s}$, 1 post-contrast PLD $= 1.5\,\mathrm{s}$, $t_L = 2\,\mathrm{s}$, $\lambda = 0.9$ and $\alpha = 0.85$.

#### Supporting information S5: systematic errors in blood $T_1$ values (in vivo data)

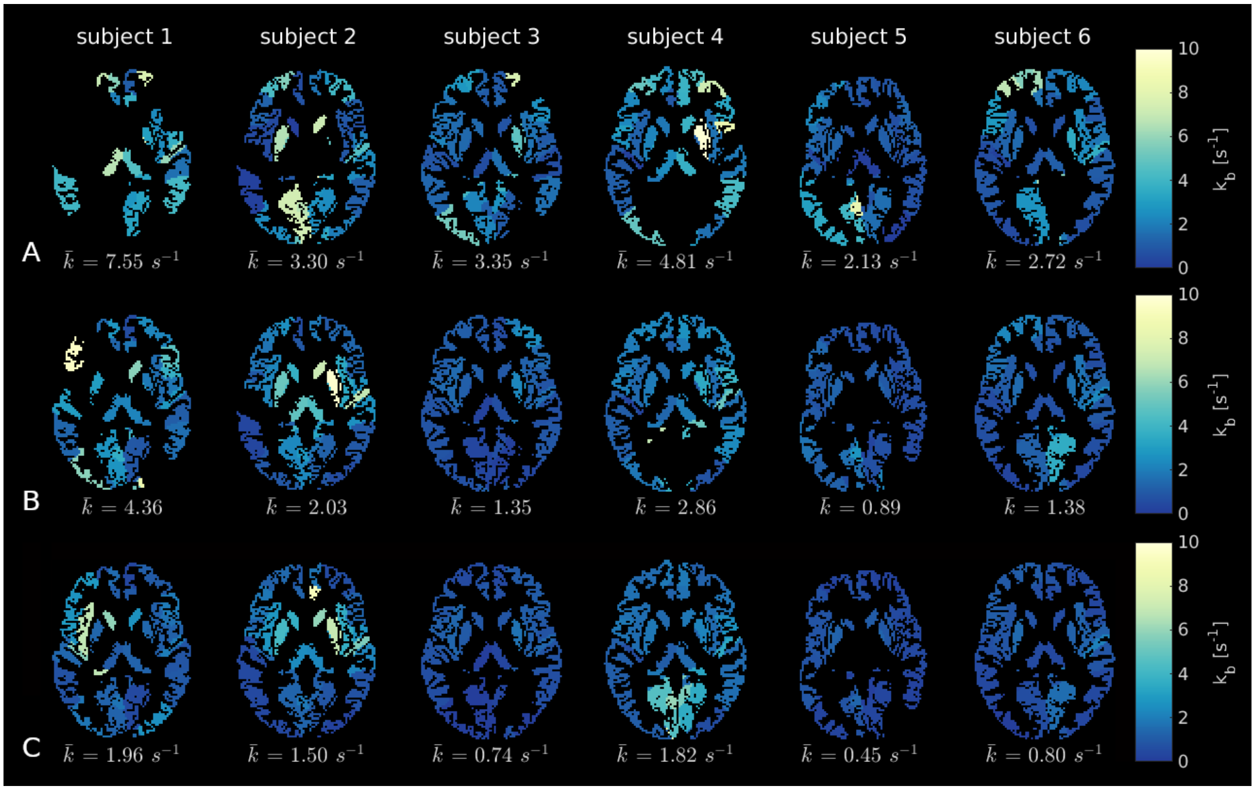


**Figure S5. Effect of systematic blood $T_1$ errors (in vivo).**
The model was fitted again to the second post-contrast (PC2) data set using adjusted $T_{1,b}$ values. The average of the ROI values, $\bar{k}$, is displayed for each volunteer. **(A)**. $T_{1,b}^{pre}$ and $T_{1,b}^{post}$ adjusted 10 % lower than measured. **(B)**. $T_{1,b}^{pre}$ and $T_{1,b}^{post}$ as measured. **(C)**. $T_{1,b}^{pre}$ and $T_{1,b}^{post}$ adjusted 10 % higher than measured.

### Supporting information S6: results from PC1

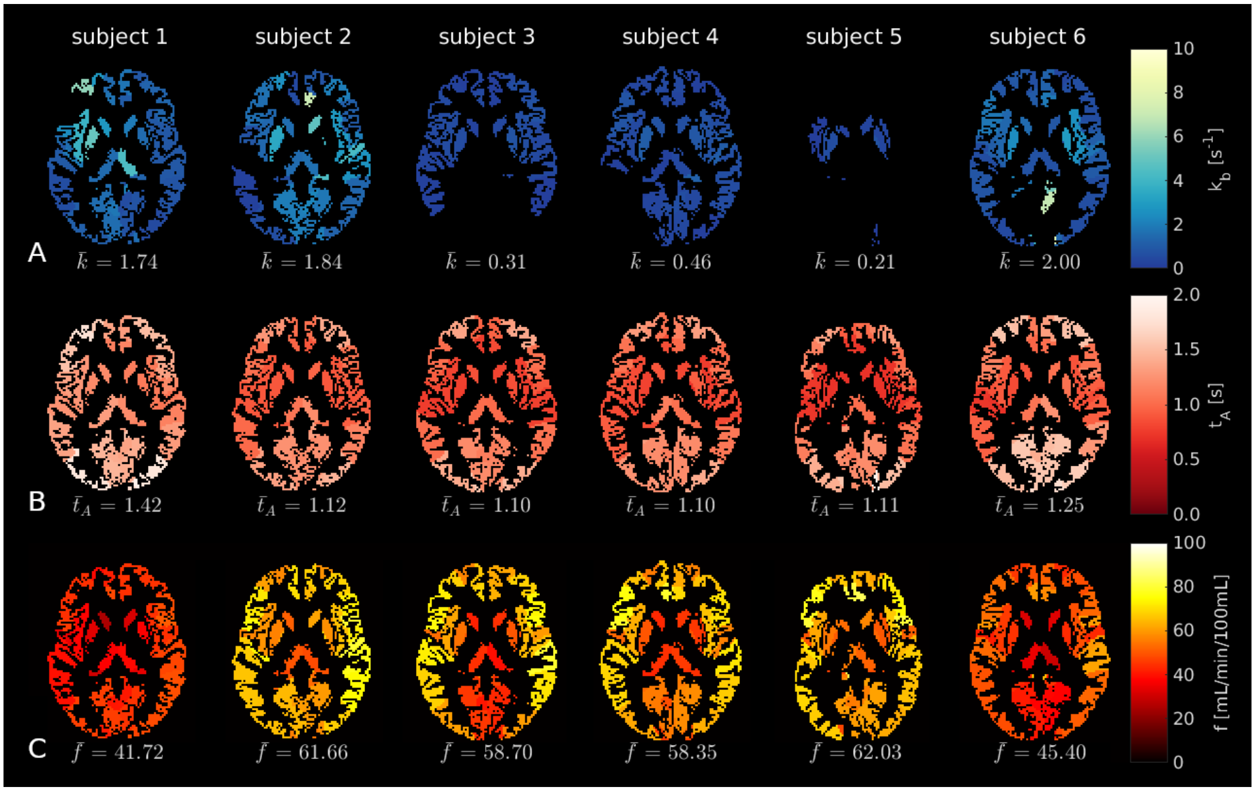


**Figure S6.1. ASL parameter maps (PC1).**

**A**. Exchange rate, $k_b$. **B**. Arterial transit time, $t_A$. **C**. Cerebral blood flow, $f$. In all maps, black voxels represent masked white matter and CSF, as well as extreme $k_b$ fits (i.e. $k_b < 0\,\mathrm{s}^{-1}$ or $k_b > 10\,\mathrm{s}^{-1}$).

**Table S6.2. ASL regional parameter fits (PC1)**.

Values are the mean and standard deviation across subjects of regional fits. GM = grey matter; FL = frontal lobe; OC = occipital lobe; PAR = parietal lobe; TLE = temporal lobe; PrcG = precentral gyrus; SFG = superior frontal gyrus; OFGsup = superior orbitofrontal gyrus; MFG = middle frontal gyrus; OFGmid = middle orbitofrontal gyrus; IFGop = inferior frontal gyrus (opercular part); IFGtr = inferior frontal gyrus (triangular part); OFGinf = inferior orbitofrontal gyrus; ROL = rolandic operculum; SMA = supplementary motor area; OLF = olfactory cortex; SFGmed = superior frontal gyrus (medial part); OFGmid = orbitofrontal gyrus (middle part); REC = rectus; INS = insula; ACC = anterior cingulate cortex; MCC = middle cingulate cortex; PCC = posterior cingulate cortex; HP = hippocampus; PHG = parahippocampal gyrus; AMYG = amygdala; CAL = calcarine; CUN = cuneus; LING = lingual gyrus; SOG = superior occipital gyrus; MOG = middle occipital gyrus; IOG = inferior occipital gyrus; FFG = fusiform gyrus; PocG = postcentral gyrus; SPG = superior parietal gyrus; IPL = inferior parietal lobule; SMG = supramarginal gyrus; ANG = angular gyrus; PRCU = precuneus; PCL = paracentral lobule; CAU = caudate; PUT = putamen; PAL = pallidum; THAL = thalamus; HES = Heschl gyrus; STG = superior temporal gyrus; TPsup = temporal pole (superior part); MTG = middle temporal gyrus; TPmid = temporal pole (middle part); ITG = inferior temporal gyrus; L = left; R = right.

| Region name | No. voxels | $f$ [ml / min / 100 ml] | $t_A$ [s] | $k_b$ [$s^{-1}$] |
|---|---|---|---|---|
| GM (L) | 55805 | $51.29 \pm 9.81$ | $1.21 \pm 0.12$ | $0.90 \pm 0.95$ |
| GM (R) | 46429 | $51.53 \pm 8.80$ | $1.20 \pm 0.13$ | $0.82 \pm 0.84$ |
| FL (L) | 12154 | $58.22 \pm 10.31$ | $1.20 \pm 0.12$ | $0.69 \pm 0.70$ |
| FL (R) | 11744 | $58.98 \pm 10.42$ | $1.20 \pm 0.15$ | $0.92 \pm 0.86$ |
| OC (L) | 6678 | $51.55 \pm 7.15$ | $1.38 \pm 0.15$ | $0.84 \pm 1.15$ |
| OC (R) | 7591 | $51.28 \pm 8.28$ | $1.37 \pm 0.17$ | $0.69 \pm 1.22$ |
| PAR (L) | 4287 | $54.03 \pm 9.38$ | $1.31 \pm 0.14$ | $0.69 \pm 0.89$ |
| PAR (R) | 4561 | $52.19 \pm 9.05$ | $1.32 \pm 0.18$ | $0.48 \pm 0.59$ |
| TLE (L) | 7046 | $59.76 \pm 10.80$ | $1.09 \pm 0.14$ | $0.71 \pm 0.69$ |
| TLE (R) | 6643 | $56.33 \pm 10.48$ | $1.04 \pm 0.13$ | $0.36 \pm 0.43$ |
| PrcG (L) | 1379 | $54.74 \pm 8.67$ | $1.34 \pm 0.24$ | $0.95 \pm 1.06$ |
| PrcG (R) | 1212 | $53.75 \pm 9.02$ | $1.32 \pm 0.13$ | $0.65 \pm 0.85$ |
| SFG (L) | 1366 | $52.42 \pm 8.86$ | $1.38 \pm 0.13$ | $1.07 \pm 1.21$ |
| SFG (R) | 1696 | $54.20 \pm 8.10$ | $1.36 \pm 0.13$ | $0.39 \pm 0.43$ |
| OFGsup (L) | 494 | $53.05 \pm 9.53$ | $1.17 \pm 0.16$ | $0.91 \pm 0.73$ |
| OFGsup (R) | 511 | $50.10 \pm 7.31$ | $1.18 \pm 0.11$ | $0.59 \pm 0.67$ |
| MFG (L) | 2142 | $60.59 \pm 11.72$ | $1.43 \pm 0.19$ | $1.43 \pm 2.09$ |
| MFG (R) | 2293 | $61.15 \pm 10.75$ | $1.40 \pm 0.11$ | $0.72 \pm 0.75$ |
| OFGmid (L) | 428 | $59.65 \pm 10.55$ | $1.28 \pm 0.16$ | $1.20 \pm 1.02$ |

| Region name | No. voxels | $f$ | $t_A$ | $k_b$ |
|---|---|---|---|---|
| OFGmid (R) | 471 | $60.45 \pm 10.49$ | $1.26 \pm 0.15$ | $1.21 \pm 1.19$ |
| IFGop (L) | 439 | $69.25 \pm 15.39$ | $1.07 \pm 0.17$ | $1.04 \pm 1.11$ |
| IFGop (R) | 615 | $69.58 \pm 11.37$ | $1.10 \pm 0.12$ | $0.60 \pm 0.63$ |
| IFGtr (L) | 1024 | $65.85 \pm 12.53$ | $1.15 \pm 0.18$ | $2.44 \pm 3.49$ |
| IFGtr (R) | 808 | $65.35 \pm 11.45$ | $1.09 \pm 0.14$ | $0.68 \pm 0.66$ |
| OFGinf (L) | 856 | $64.03 \pm 12.86$ | $1.05 \pm 0.17$ | $2.41 \pm 3.51$ |
| OFGinf (R) | 768 | $61.14 \pm 9.75$ | $0.94 \pm 0.20$ | $1.01 \pm 0.76$ |
| ROL (L) | 514 | $47.87 \pm 7.93$ | $0.93 \pm 0.19$ | $0.75 \pm 0.69$ |
| ROL (R) | 689 | $54.71 \pm 11.23$ | $0.97 \pm 0.11$ | $0.88 \pm 0.64$ |
| SMA (L) | 905 | $54.88 \pm 10.39$ | $1.16 \pm 0.16$ | $0.54 \pm 0.47$ |
| SMA (R) | 1016 | $52.06 \pm 8.11$ | $1.18 \pm 0.19$ | $0.90 \pm 0.94$ |
| OLF (L) | 187 | $55.23 \pm 11.35$ | $0.84 \pm 0.17$ | $1.48 \pm 1.54$ |
| OLF (R) | 189 | $52.79 \pm 10.15$ | $0.89 \pm 0.14$ | $1.50 \pm 1.54$ |
| SFGmed (L) | 1162 | $60.22 \pm 11.89$ | $1.07 \pm 0.14$ | $0.53 \pm 1.07$ |
| SFGmed (R) | 983 | $58.58 \pm 8.63$ | $1.12 \pm 0.14$ | $0.71 \pm 0.67$ |
| OFGmed (L) | 370 | $70.95 \pm 14.19$ | $1.00 \pm 0.18$ | $0.67 \pm 0.56$ |
| OFGmed (R) | 463 | $66.54 \pm 9.63$ | $1.00 \pm 0.15$ | $1.13 \pm 1.63$ |
| REC (L) | 478 | $67.90 \pm 12.05$ | $0.93 \pm 0.16$ | $0.48 \pm 0.52$ |
| REC (R) | 440 | $59.51 \pm 8.32$ | $0.96 \pm 0.13$ | $0.78 \pm 1.06$ |
| INS (L) | 1243 | $54.92 \pm 9.73$ | $0.91 \pm 0.18$ | $1.28 \pm 1.28$ |
| INS (R) | 1097 | $59.46 \pm 10.43$ | $0.85 \pm 0.13$ | $1.24 \pm 1.08$ |
| ACC (L) | 812 | $68.02 \pm 13.40$ | $0.90 \pm 0.13$ | $0.67 \pm 0.90$ |
| ACC (R) | 753 | $63.87 \pm 11.87$ | $0.94 \pm 0.15$ | $1.74 \pm 2.52$ |
| MCC (L) | 1095 | $58.77 \pm 9.66$ | $1.08 \pm 0.10$ | $1.08 \pm 1.23$ |
| MCC (R) | 1292 | $60.13 \pm 11.17$ | $1.07 \pm 0.17$ | $1.84 \pm 2.16$ |
| PCC (L) | 192 | $66.03 \pm 11.41$ | $1.19 \pm 0.08$ | $2.21 \pm 3.77$ |
| PCC (R) | 110 | $63.30 \pm 10.97$ | $1.24 \pm 0.13$ | $2.20 \pm 4.68$ |
| HP (L) | 614 | $40.63 \pm 11.61$ | $0.92 \pm 0.25$ | $0.80 \pm 3.58$ |
| HP (R) | 530 | $41.72 \pm 7.83$ | $0.99 \pm 0.13$ | $1.38 \pm 1.48$ |
| PHG (L) | 555 | $43.62 \pm 9.02$ | $0.93 \pm 0.16$ | $1.04 \pm 1.32$ |
| PHG (R) | 698 | $44.33 \pm 8.63$ | $0.94 \pm 0.11$ | $0.99 \pm 1.50$ |
| AMYG (L) | 174 | $42.24 \pm 9.16$ | $0.90 \pm 0.18$ | $1.71 \pm 2.19$ |
| AMYG (R) | 187 | $42.74 \pm 6.75$ | $0.89 \pm 0.15$ | $2.67 \pm 3.05$ |
| CAL (L) | 1266 | $52.19 \pm 10.28$ | $1.31 \pm 0.20$ | $2.22 \pm 3.78$ |
| CAL (R) | 1010 | $50.63 \pm 10.03$ | $1.30 \pm 0.17$ | $2.06 \pm 4.71$ |
| CUN (L) | 715 | $47.04 \pm 6.92$ | $1.37 \pm 0.14$ | $1.18 \pm 3.66$ |
| CUN (R) | 723 | $49.65 \pm 7.01$ | $1.43 \pm 0.20$ | $1.72 \pm 2.96$ |
| LING (L) | 1253 | $47.05 \pm 8.79$ | $1.22 \pm 0.22$ | $2.31 \pm 4.12$ |

| Region name | No. voxels | $f$ | $t_A$ | $k_b$ |
|---|---|---|---|---|
| LING (R) | 1296 | $49.17 \pm 9.87$ | $1.26 \pm 0.14$ | $1.62 \pm 4.06$ |
| SOG (L) | 549 | $45.56 \pm 6.26$ | $1.56 \pm 0.16$ | $0.28 \pm 2.58$ |
| SOG (R) | 576 | $53.22 \pm 8.08$ | $1.66 \pm 0.24$ | $0.83 \pm 0.88$ |
| MOG (L) | 1716 | $58.47 \pm 8.97$ | $1.45 \pm 0.21$ | $0.24 \pm 0.70$ |
| MOG (R) | 1035 | $58.76 \pm 7.95$ | $1.52 \pm 0.14$ | $0.28 \pm 0.57$ |
| IOG (L) | 542 | $57.00 \pm 11.60$ | $1.44 \pm 0.21$ | $0.32 \pm 0.62$ |
| IOG (R) | 495 | $55.36 \pm 10.54$ | $1.46 \pm 0.17$ | $0.17 \pm 0.47$ |
| FFG (L) | 1562 | $39.61 \pm 7.98$ | $1.18 \pm 0.21$ | $1.23 \pm 1.34$ |
| FFG (R) | 1635 | $38.92 \pm 8.66$ | $1.20 \pm 0.07$ | $1.35 \pm 1.78$ |
| PoCG (L) | 1473 | $52.10 \pm 12.63$ | $1.30 \pm 0.19$ | $0.53 \pm 1.55$ |
| PoCG (R) | 1341 | $48.30 \pm 8.02$ | $1.31 \pm 0.11$ | $0.51 \pm 0.69$ |
| SPG (L) | 704 | $43.17 \pm 5.88$ | $1.58 \pm 0.21$ | $0.82 \pm 1.47$ |
| SPG (R) | 620 | $42.77 \pm 7.24$ | $1.60 \pm 0.19$ | $0.79 \pm 0.82$ |
| IPL (L) | 1165 | $51.80 \pm 11.24$ | $1.35 \pm 0.25$ | $0.68 \pm 0.97$ |
| IPL (R) | 583 | $54.35 \pm 9.02$ | $1.37 \pm 0.14$ | $0.75 \pm 0.88$ |
| SMG (L) | 599 | $57.87 \pm 11.70$ | $1.11 \pm 0.20$ | $0.39 \pm 0.50$ |
| SMG (R) | 921 | $62.10 \pm 11.52$ | $1.10 \pm 0.11$ | $0.82 \pm 0.87$ |
| ANG (L) | 621 | $58.81 \pm 10.25$ | $1.29 \pm 0.15$ | $0.46 \pm 0.69$ |
| ANG (R) | 822 | $59.76 \pm 8.74$ | $1.31 \pm 0.11$ | $0.73 \pm 1.07$ |
| PRCU (L) | 1551 | $48.65 \pm 5.14$ | $1.32 \pm 0.10$ | $0.76 \pm 1.11$ |
| PRCU (R) | 1544 | $50.53 \pm 6.80$ | $1.34 \pm 0.14$ | $1.46 \pm 2.01$ |
| PCL (L) | 420 | $41.54 \pm 5.21$ | $1.29 \pm 0.11$ | $0.46 \pm 1.92$ |
| PCL (R) | 287 | $43.22 \pm 6.65$ | $1.34 \pm 0.15$ | $0.51 \pm 0.71$ |
| CAU (L) | 545 | $41.15 \pm 10.01$ | $0.91 \pm 0.24$ | $1.19 \pm 0.92$ |
| CAU (R) | 584 | $45.39 \pm 12.31$ | $0.91 \pm 0.15$ | $1.50 \pm 1.59$ |
| PUT (L) | 674 | $48.80 \pm 10.08$ | $0.87 \pm 0.17$ | $2.18 \pm 1.70$ |
| PUT (R) | 690 | $47.33 \pm 6.74$ | $0.92 \pm 0.13$ | $2.74 \pm 3.35$ |
| PAL (L) | 71 | $49.80 \pm 8.53$ | $0.89 \pm 0.20$ | $2.21 \pm 2.58$ |
| PAL (R) | 67 | $50.76 \pm 9.73$ | $0.85 \pm 0.13$ | $1.52 \pm 2.62$ |
| THAL (L) | 357 | $44.20 \pm 7.20$ | $1.10 \pm 0.11$ | $0.54 \pm 0.79$ |
| THAL (R) | 399 | $42.25 \pm 7.78$ | $1.12 \pm 0.11$ | $1.03 \pm 1.67$ |
| HES (L) | 136 | $62.60 \pm 17.52$ | $0.84 \pm 0.24$ | $0.55 \pm 0.74$ |
| HES (R) | 139 | $62.10 \pm 10.66$ | $0.86 \pm 0.12$ | $1.40 \pm 1.41$ |
| STG (L) | 1142 | $61.08 \pm 11.45$ | $0.87 \pm 0.18$ | $0.33 \pm 0.65$ |
| STG (R) | 1474 | $67.33 \pm 11.97$ | $0.91 \pm 0.14$ | $0.93 \pm 0.84$ |
| TPsup (L) | 451 | $57.33 \pm 11.68$ | $0.85 \pm 0.13$ | $0.55 \pm 0.44$ |
| TPsup (R) | 515 | $57.92 \pm 10.96$ | $0.87 \pm 0.18$ | $1.16 \pm 1.22$ |
| MTG (L) | 2696 | $60.07 \pm 12.00$ | $1.06 \pm 0.15$ | $0.24 \pm 0.79$ |

| Region name | No. voxels | $f$ | $t_A$ | $k_b$ |
|---|---|---|---|---|
| MTG (R) | 2427 | $63.24 \pm 10.77$ | $1.15 \pm 0.13$ | $0.52 \pm 0.54$ |
| TPmid (L) | 342 | $50.72 \pm 9.68$ | $1.03 \pm 0.13$ | $1.40 \pm 1.55$ |
| TPmid (R) | 482 | $49.61 \pm 6.32$ | $1.00 \pm 0.16$ | $1.23 \pm 1.35$ |
| ITG (L) | 1876 | $45.22 \pm 6.99$ | $1.18 \pm 0.13$ | $0.51 \pm 0.72$ |
| ITG (R) | 2009 | $52.48 \pm 9.77$ | $1.23 \pm 0.10$ | $0.64 \pm 0.67$ |
